# CryoEM reveals the dynamic conformational landscape and C-terminal gating mechanism of Mycobacterial Class Ib ribonucleotide reductase

**DOI:** 10.64898/2026.06.02.729529

**Authors:** Lumbini R. Yadav, Pratibha Tiwari, Janesh Kumar, Shekhar C. Mande

## Abstract

Ribonucleotide reductases (RNRs) are essential enzymes that catalyze the reduction of ribonucleoside diphosphates to deoxyribonucleoside diphosphates and function as multi-subunit, asymmetric heterotetrameric (α₂β₂) complexes. Despite their central role in DNA synthesis, the mechanism of subunit association and regulation within this asymmetric assembly has remained incompletely understood. Here, we employed cryo–electron microscopy (cryo-EM) to investigate the apo state assembly and dynamics of a class Ib RNR complex from *Mycobacterium thermoresistibile*. The 3.8 Å structure provides a detailed architecture of the complex, including order-disorder transition in the C-terminal tail of the β subunit, which is critical for long-range radical transfer. The alternative β-hairpin conformation in the α subunit suggests its role in stabilizing the α₂β₂ interaction. Structural analysis of the α₂β₂ complex identified seven distinct conformational states, highlighting substantial heterogeneity and variability in α–β subunit association. The observed structural heterogeneity supports a model for apo-state dynamics and suggests β-subunit movements toward the α subunit that may facilitate productive αβ and potential α′β′ interactions. Complementary thermodynamic analyses further support a model in which conformational flexibility is central to α–β subunit recognition and stabilization. Together, these findings illuminate the dynamic conformational landscape that governs subunit association and radical transfer in class Ib RNRs. Given the importance of mycobacteria as pathogens of significant importance in humans and animals, this first structural characterization of a mycobacterial class Ib RNR provides a foundation for future structure-based inhibitor design.

**Significance Statement:** RNRs are essential for DNA synthesis in all life forms. Here, we present cryo-EM structures of the ligand-free mycobacterial α₂β₂ complex, capturing multiple modes of asymmetric subunit association. These structures show that the enzyme samples multiple conformations that likely correspond to different activity states, potentially associated with processes such as DNA repair or replication. Notably, this flexibility arises intrinsically from subunit interactions, even in the absence of substrates or regulatory nucleotides. A 3.8 Å structure reveals ordering in the β-subunit C-terminal tail critical for long-range radical transfer, while an alternative β-hairpin in the α subunit may stabilize the complex. These results provide a structural framework for RNR function and inform future drug design.

## 1. Introduction

A balanced supply of deoxyribonucleotides plays a crucial role in maintaining the genomic stability and the fidelity during DNA replication and repair processes. Ribonucleotide reductase complex (RNR) maintains this balance through well-coordinated allosteric regulatory mechanisms^1,2^. RNR catalyzes the reduction of ribonucleoside diphosphates to deoxyribonucleoside diphosphates by abstraction of the 2’ hydroxyl group from the ribose sugar through a radical transfer (RT) mechanism ^3,4^. Based on the cofactor required, oxygen-dependence, radical initiation pathway, and other factors, RNRs are classified into three major classes: Class I (a-e), Class II, and Class III ^5,6,7^. *Mycobacterium tuberculosis* (Mtb) possesses class Ib and class II RNRs. Class Ib RNR is encoded by a gene cluster (*nrdHIEF2*) essential for Mtb growth. Moreover, other genes for the NrdF/β subunits, namely *nrdF1* (class Ib-like) and *nrdB* (class Ic-like), are also present ^8,9^. The small subunits encoded by *nrdF1*, *nrdB* as well as *nrdZ* genes are unable to substitute for the *nrdF2* encoded small subunit, proving its indispensability ^10,11^. Studying the structure and function of these RNRs is thus important, as these are promising targets against tuberculosis ^12,13,14^.

Class Ia RNR found in humans or in bacteria such as *Escherichia coli* is one of the most comprehensively studied classes of ribonucleotide reductase and has provided much of the information available about the RNR mechanism. Extensive spectroscopic and structural analyses have been performed on class Ia RNRs, revealing the details of structures and intricate mechanism of radical transfer ^15,16,17,18,19^. Class Ib RNR, found exclusively in prokaryotes and mostly in pathogenic bacteria, have several characteristic features that distinguish them from class Ia RNR. For example, the cone domain at the N-terminus that is present in class Ia RNR is absent in class Ib RNR ^1,20,21^, class Ia RNR contains a diferric tyrosyl radical cofactor, whereas class Ib RNR can be activated by either iron or manganese, ^22^ and so on. The two subunits of the class Ib RNR enzyme, NrdE and NrdF, are more commonly called α and β subunits, respectively.

The prototypical class I RNR is an asymmetric tetramer composed of two homodimeric subunits, α and β. The α subunit contains catalytic site for substrate reduction, an allosteric site for regulation, and redox-active thiols/disulphides essential for catalysis. The β subunit houses oxygen-linked metal cofactor site, where the radical required for catalysis is generated and transfer is initiated. In class Ib RNR, the presence of manganese at the metallocofactor site requires the accessory subunit NrdI because manganese alone cannot form a tyrosyl radical in the presence of atmospheric oxygen. NrdI, a flavodoxin-like protein, facilitates radical generation by producing oxidants through oxygen reduction, which are then used to assemble the radical at the metallocofactor site in the β-subunit ^23,24,22^. Our recently determined structure of the Mycobacterial NrdF₂:NrdI complex reveals a detailed mechanism for Tyr-radical formation, including an oxygen tunnel leading to the FMN site of NrdI, where oxygen is reduced to superoxide. The superoxide radical is then transferred through a tunnel across the NrdI and NrdF subunits to the metal-binding site. Once there, the activated metal oxidizes tyrosine to form tyrosyl radical ^23,25,26^. Once the tyrosyl radical is formed, the proton-coupled electron transport (PCET) pathway initiates from the β subunit through residues Y122, W48, and Y356. Eventually, it lands in the α subunit at residues Y731, Y730, and C439 at a distance of 35-40 Å away^27,28^. Several studies through site-directed mutagenesis along with the incorporation of unnatural amino acids and formation of semisynthetic β subunit using expressed protein ligation method have confirmed the role of residues involved in PCET in both the α_2_ and β_2_ subunits ^29,30,31,32,33^.

Given the asymmetric nature of the α_2_β_2_ complex, inter-subunit interactions lead to only half of the tyrosines (Y122) in the β subunit and half of the redox-active cysteine residues (C439) in the α subunit being catalytically active under a single-turnover conditions ^34,35,36^. This phenomenon, known as half-site reactivity, has been demonstrated through structural studies of RNRs of *Salmonella typhimurium* and *E. coli*. Similarly, the PCET has been well documented in class Ia RNR ^29,30,31,32,33^. Our structural analysis of the NrdE (α-subunit) dimer suggests that half-site reactivity may not solely arise from asymmetry in the α_2_β_2_ complex, but could also be an intrinsic property of the α-subunit itself ^37^. However, structural information for class Ib RNRs remains limited. The only class Ib X-ray structure reported at 4 Å represents an intermediate state ^38^. Recently, the cryo-EM structure of the Class Ib α₂β₂ complex from *B. subtilis* revealed wide conformational landscape, ranging from open to compact states, including PCET-competent conformation ^39^. In contrast, the *E. coli* class Ia RNR structure was captured in a trapped state using doubly substituted β_2_ variant (E52Q/(2,3,5)-trifluorotyrosine122), and its cryo-EM structure was the first to delineate the complete free radical pathway by resolving the entire C-terminal tail of the β subunit ^16^.

This study aims to elucidate the intrinsic, ligand-free conformational landscape of asymmetric α_2_β_2_ complexes in class Ib ribonucleotide reductases. We focus specifically on Mycobacterial class Ib RNR, encoded by the *nrdE* and *nrdF* genes, in ligand-free state, with the broader goal of expanding structural insights into Mycobacterial proteins^12^. By integrating Cryo-EM data with thermodynamic analyses, we characterize the molecular assembly, subunit interactions, and the probable conformational transitions associated with radical transfer between subunits in class Ib RNR. The 3.8 Å cryo-EM structure of the α_2_β_2_ complex of *Mycobacterium thermoresistibile* (Mth) reveals an overall asymmetric architecture, as expected, but with notable differences in its mode of assembly compared to previously reported RNR structures ^39^. Conformational variability analysis indicates that the flexible C-terminal of the β subunit becomes progressively ordered as it approaches the α subunit during radical transfer. Notably, we identify as many as seven distinct conformational states within the α_2_β_2_ complex. Both local and global conformational variability in the subunits suggests a coordinated mechanism of α’ and β’ interaction. Collectively, these findings provide a dynamic structural framework of the α_2_β_2_ complex, offering insights into the stepwise association of the two dimers and the conformational transitions that likely accompany long-range free radical transfer.

## 2. Results

Ribonucleotide reductases catalyze the essential reduction of ribonucleosides to deoxyribonucleosides, a key step in DNA synthesis and repair. In class Ib RNRs, the subunits NrdE (α), NrdF (β), and NrdI work in tandem to facilitate this process. In this study, we used cryo-EM single-particle analysis to investigate the ligand-free assembly of the RNR complex and gain mechanistic insights into its function and dynamic behaviour.

### 2.1. Cryo-EM analysis of the ternary NrdEFI complex

The final set of selected particles (#716K) derived from a large data set (#6935 micrographs) of the ternary complex was subjected to *ab initio* reconstruction into twelve classes **(Supplementary Figures 1A and 1B).** Analysis of the 12 reconstructed maps revealed two distinct groups. The first group (a), comprising six classes, exhibited poor inter-subunit interface density and was classified as dissociated states. These maps showed well-defined density for the α subunit but weak or diffuse density for the β subunit. The narrow α–β interface suggests a weak or transient interaction; therefore, these classes were excluded from further analysis. Notably, these excluded maps closely resemble the overall architecture of class Ib α_2_β_2_ structure reported for *B. subtilis* ^39^. The second group (b) consisted of five classes with well-defined inter-subunit interface density. These class with intact particles, designated as conformers A, C, E, F, and G, each containing ∼10,000–17,000 particles, were selected for further reconstruction **(Supplementary Figure 1D).** All selected classes exhibited a clearly resolved α–β interface, indicative of stable subunit association. However, the density for NrdI subunit in this ternary complex is detectable but very poor. The conformational heterogeneity, preferential orientation and compositional variability resulting from ternary complex dissociation on the grid, limited the resolution of the resulting maps. To address these issues of heterogeneity and resolution, we shifted our approach to studying a more stable binary α_2_β_2_ protein complex.

2.2. **Cryo-EM analysis of the α₂β₂ binary complex**

The α_2_β_2_ protein complex was obtained by co-expression and co-purification using affinity and size exclusion chromatography **(Supplementary Figure 2).** To further minimize dissociation and heterogeneity, α_2_β_2_ protein complex was crosslinked with glutaraldehyde. Crosslinking of the α_2_β_2_ complex significantly improved particle uniformity, enriching the sample with one major population **(Supplementary Figure 3).** The structure of this enriched conformer of the α_2_β_2_ protein complex (conformer D) was obtained at a global resolution of 3.8 Å **(Supplementary Figure 3C, 3E, and 3F).** In addition to this, another conformer (conformer B) was determined at a lower resolution of 6.1 Å, while particles from other classes were not refined due to their poor map quality **(Supplementary Figure 3D and 3E).** Interestingly, crosslinking did not alter the mode of α–β interaction, as a similar mode of interaction was observed between the subunits even in the absence of crosslinking agent, allowing us to use the better-resolved maps of the ternary complex to interpret α–β conformational variability **(Supplementary Figure 1D).**

### 2.3. Cryo-EM structure reveals seven distinct conformations

Across ternary and binary complexes, seven distinct conformational states were observed in the cryo-EM maps, named as conformers A to G **(Figure 1)**. The conformers B and D are obtained from the cross-linked α_2_β_2_ binary complex **(Supplementary Figure 3)** and conformers A, C, E, F, and G are obtained from the ternary complex (**Supplementary Figure 1)**. These conformers can be clustered into three major groups based on the α–β interaction pattern: 1. Extended conformation (conformers A and B) **(Figure 1B),** 2. Intermediate conformation (conformer C, D, F, G) **(Figure 1C)**, and 3. Compact conformation (conformer E) **(Figure 1D).** We modelled and attempted refinement for α_2_ and β_2_ subunits in all the conformers, but final refinement and submission was done only for conformer D **(Figure 1C).** Only the resolution of Conformer D was suitable for side-chain modelling. Conformers A, B, C, F and G were only used for domain-level analysis. **Table 1** summarizes map details for all the conformers with validation statistics of the cryo-EM structure determined for conformer D. In subsequent sections, the interacting chains are labelled as α and β, whereas the non-interacting chains are labelled as α’ and β’ for the two subunits.

**Figure 1:**
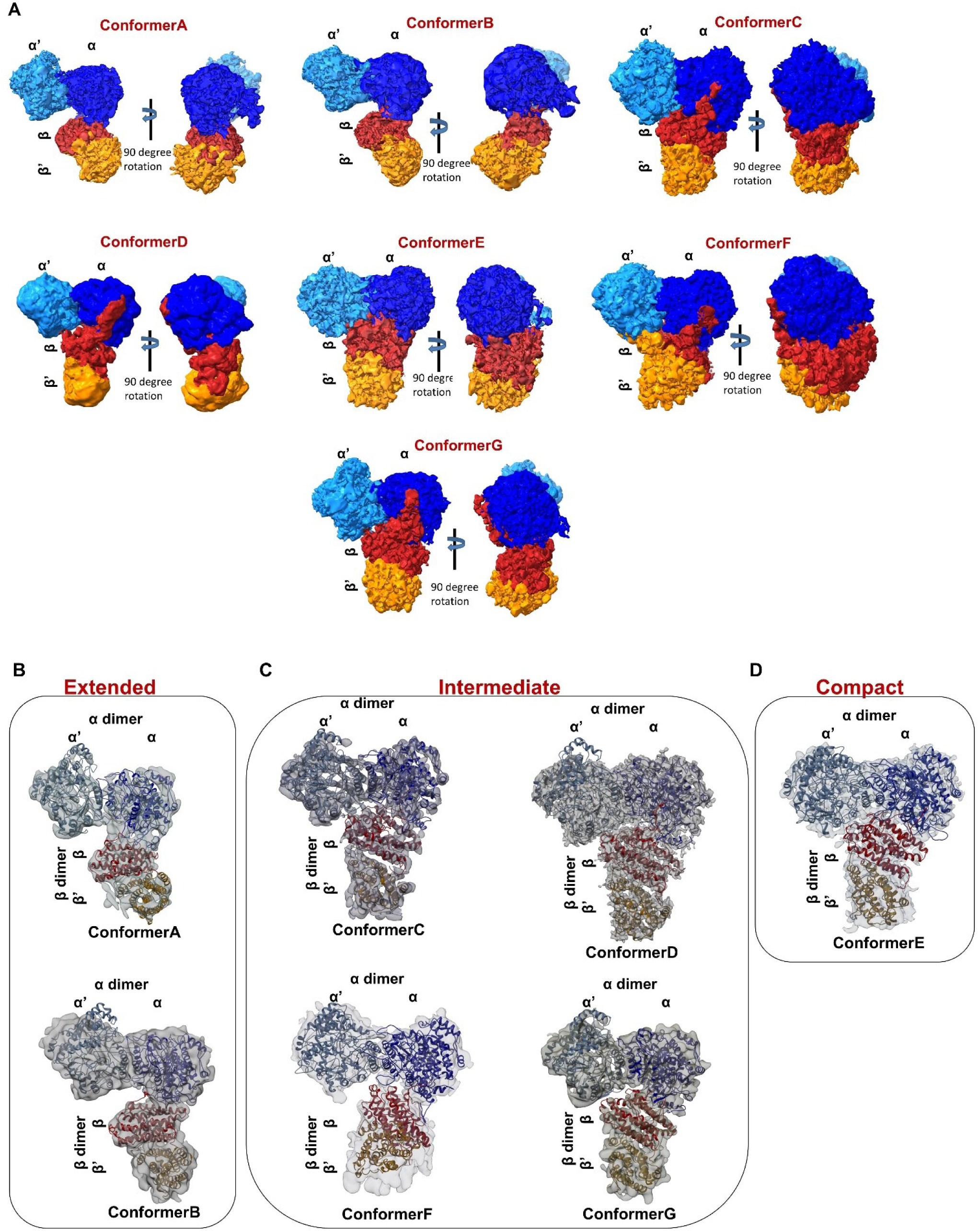
Cryo-EM maps and Map model overlay of different conformational states. **A.** Unsharpened cryo-EM maps of different conformers i.e. Conformer A, Conformer B, Conformer C, Conformer D, Conformer E, Conformer F, and Conformer G are presented in front and side views to highlight variability. The maps of different conformational states are labelled as α (blue), α’ (light blue), β (red) and β’(orange). α and β are interacting chains of NrdE and NrdF in the asymmetric complex, and α’ and β’ are non-interacting parts of the dimer. For uniformity of visualization, orientation of the α_2_ dimer is maintained in the same frame for all conformers. The sharpened maps, with the models overlaid, display various conformational states that were clustered based on their similarities, as shown. **B.** Extended conformation (conformers A and B), **C**. Intermediate conformation (conformer C, D, G) and tilted intermediate conformer (conformer F), and **D.** Compact conformation (conformer E). Side-chain modelling was only feasible for conformer D.

**Table 1:** Cryo-EM Data collection and processing details: Cryo-EM data collection and map details of conformer A-C and E-G with refinement and validation statistics of conformer D.

| Data collection and processing | Conformer A | Conformer B | Conformer C | Conformer D | Conformer E | Conformer F | Conformer G |
| --- | --- | --- | --- | --- | --- | --- | --- |
| Microscope |  |  |  | Titan Krios G3 |  |  |  |
| Voltage (kV) | 300 |  | 300 | 300 | 300 | 300 | 300 |
| No of micrographs | 6935 | 831 | 6935 | 831 | 6935 | 6935 | 6935 |
| Detector | K2 | K2 | K2 | K2 | K2 | K2 | K2 |
| Exposure time (s) | 5 | 8 | 5 | 8 | 5 | 5 | 5 |
| Electron dose (e-/Å2) | 42.02 | 47.96 | 42.02 | 47.96 | 42.02 | 42.02 | 42.02 |
| Pixel size (Å) | 1.052 | 1.065 | 1.052 | 1.065 | 1.052 | 1.052 | 1.052 |
| No of particles in final map | 10664 | 14633 | 14107 | 112390 | 15807 | 17261 | 16462 |
| Map resolution (Å) | 7.47 | 6.15 | 4.64 | 3.8 | 4.31 | 4.57 | 4.53 |
| FSC threshold | 0.143 | 0.143 | 0.143 | 0.143 | 0.143 | 0.143 | 0.143 |
| Map resolution range (Å) | 7.5-15 | 6.0-15 | 4.6-10 | 3.3-10 | 4.3-15 | 4.5-15 | 4.5-15 |
| Refinement | Refinement |  |  |  |  |  |  |
| Initial model used (PDB code) | 2bq1 (NrdE dimer) |  | 8J4Y (NrdF dimer) |  |  |  |  |
| Resolution used in refinement |  |  |  | 3.8 |  |  |  |
| Map vs model FSC |  |  |  |  |  |  |  |
| CC (mask) |  |  |  | 0.6 |  |  |  |
| CC (box) |  |  |  | 0.73 |  |  |  |
| CC (peaks) |  |  |  | 0.55 |  |  |  |
| CC (volume) |  |  |  | 0.59 |  |  |  |
| CC (ligands) |  |  |  | 0.73 |  |  |  |
| B-factor (Å2) |  |  |  |  |  |  |  |
| Total |  |  |  |  |  |  |  |
| Protein |  |  |  | 104.3 |  |  |  |
| Ligand |  |  |  | 130.7 |  |  |  |
| Root mean square deviations |  |  |  |  |  |  |  |
| Bond lengths (Å) |  |  |  | 0.002 |  |  |  |
| Bond angles (°) |  |  |  | 0.6 |  |  |  |
| Validation |  |  |  |  |  |  |  |
| Molprobity score |  |  |  | 2.2 |  |  |  |
| Poor Rotamers (%) |  |  |  | 1.25 |  |  |  |
| Clashscore |  |  |  | 10.9 |  |  |  |
| Ramachandran plot (%) |  |  |  |  |  |  |  |
| Favored |  |  |  | 88.65 |  |  |  |
| Allowed |  |  |  | 10.81 |  |  |  |
| Outliers |  |  |  | 0.55 |  |  |  |

Across all maps, the α subunits consistently exhibited better resolution than the β subunits. **(Supplementary Figure 1 and 3).** Backbone chain density for most regions of the α subunits could be traced, however, the N-terminal region exhibited fragmented density. Interestingly, the α chain engaged in interaction with the β chain retained much of its N-terminal density, suggesting its stabilization upon α–β association. Notably, in Conformer F, the density in the β subunit is particularly poorly resolved, rendering backbone tracing difficult over much of the region **(Supplementary Figure 1D and Figure 1C).** Despite this, in all the conformers, both the α_2_ and the β_2_ subunits retain dimeric architecture consistent with previously reported structures. The α₂ subunit adopts a canonical dimeric architecture, with only minor conformational variations arising due to domain motions. Importantly it is this α_2_ canonical form that engages with the β_2_ subunit to assemble the α_2_β_2_ complex.

### 2.4. Global conformational changes across conformers

**Supplementary Figures 4A-4G** highlight the broad variations in assembly observed in the α and β subunits of α_2_β_2_ complex. As the mode of association between the α and β dimers differs significantly, we assessed rotations in the heterodimer by first superposing the α chain to bring them into single frame of reference for all the seven conformers (and that of *S. typhimurium*) and then calculating transformations required to superpose β-β’ chains with respect to this frame of reference. Taking the α subunit of conformer D as the template and comparing it with six other conformers showed rotations of β dimers in values ranging from 62.6° (conformer F) to 1.5° (conformer C). **Supplementary Table 3** illustrates the rotation of the β and β’ chain relative to the α chain. However, the *S. typhimurium* structure required a large rotation of 83.5° and 103° for β and β’, respectively. Thus, significant conformational changes are seen in the juxtapositions of β dimer relative to the α subunit **(Figure 2).** The major transition observed in different structures of conformers A to G, as we hypothesize, is demonstrated in **Supplementary Video 1.**

**Figure 2:**
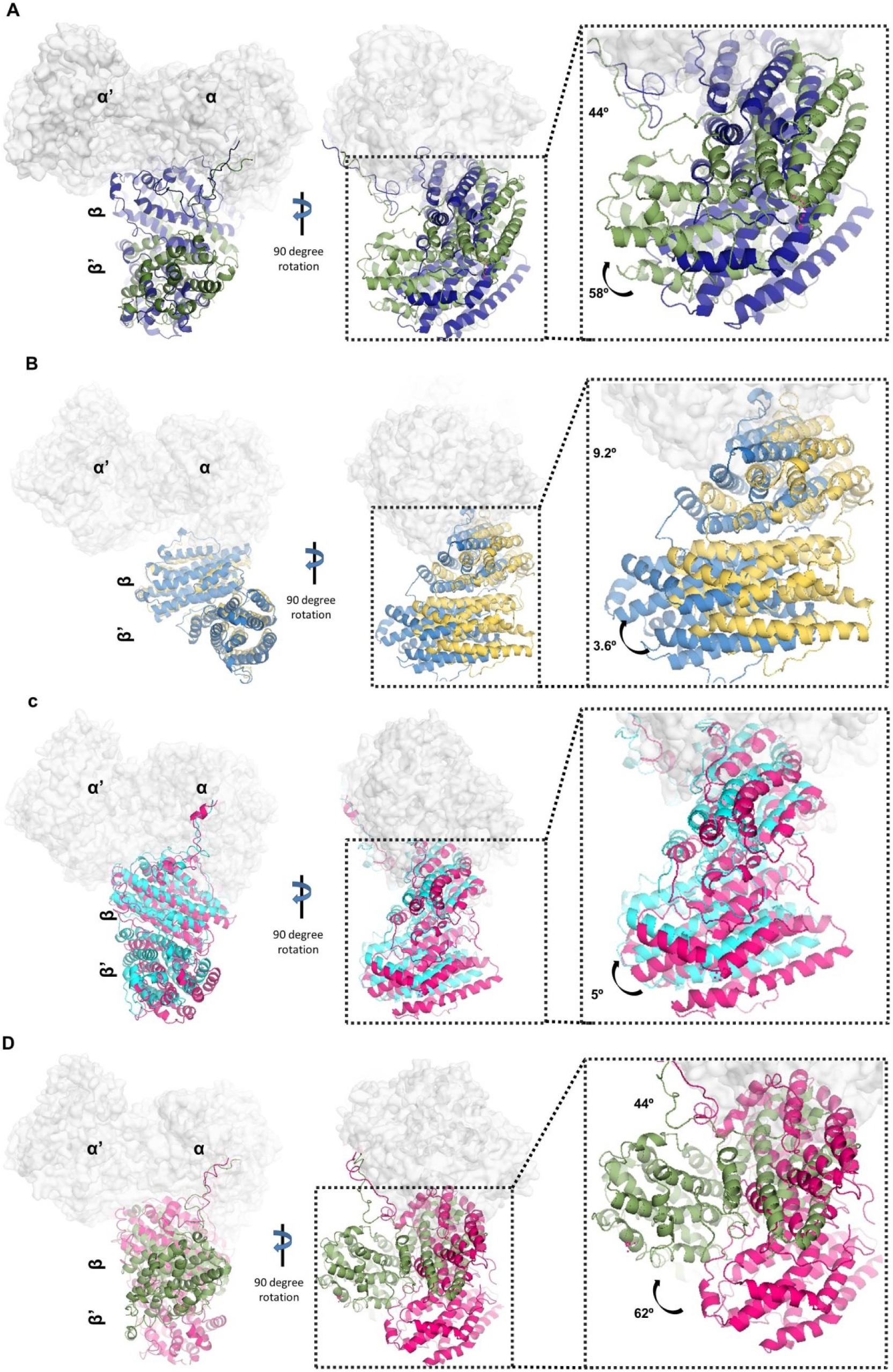
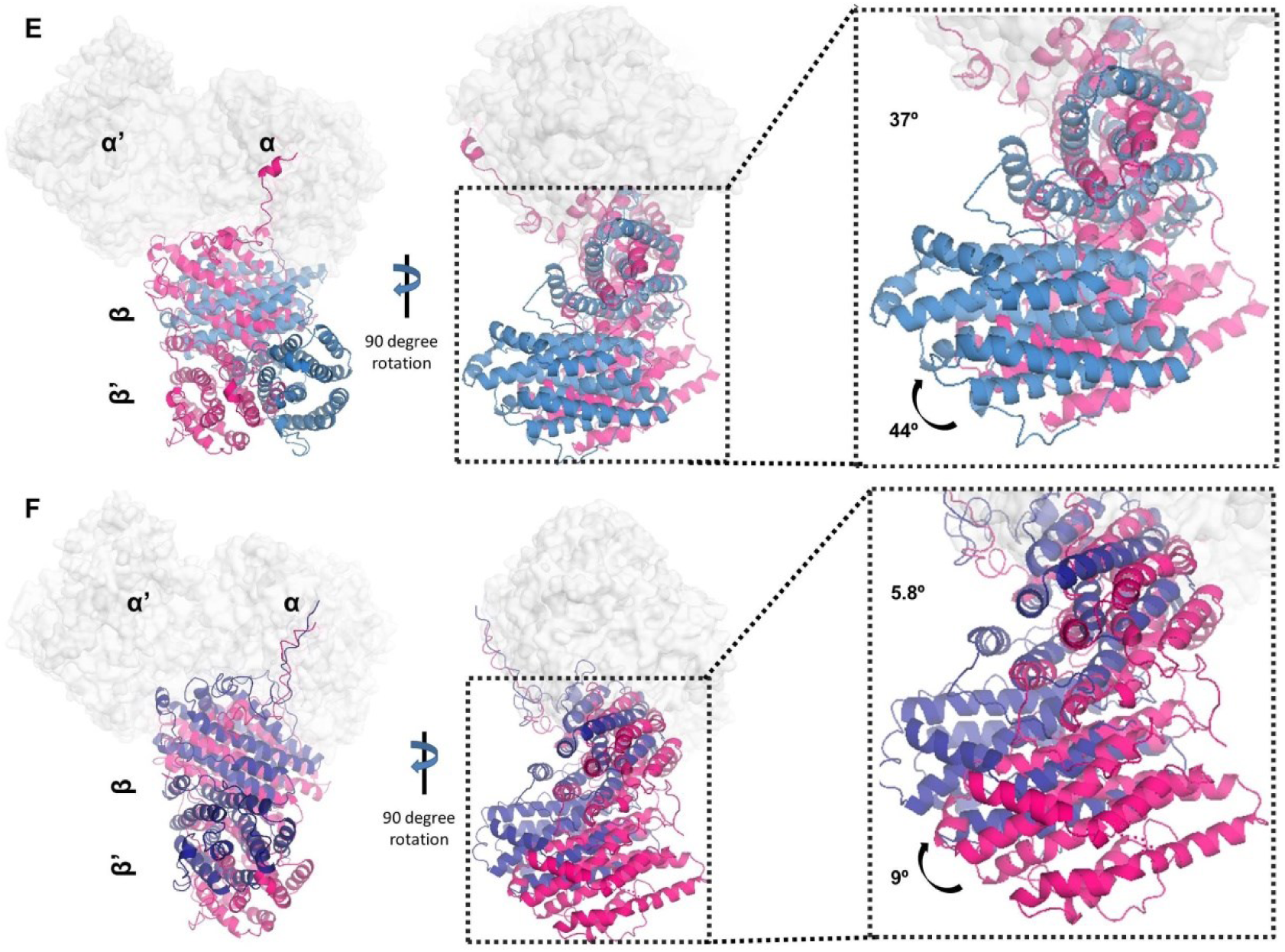
Global conformational change observed in conformers. The different conformers of α_2_β_2_ complex are superposed, and global change is highlighted in different conformers as seen in **A.** Conformer E (blue) and Conformer F (smudge-green), **B.** Conformer B (sky blue): Conformer A (yellow), **C.** Conformer D (pink): Conformer C (cyan), **D.** Conformer D (pink): Conformer F (green), **E.** Conformer D (pink): Conformer B (sky blue), **F.** Conformer D (pink): Conformer E (blue). Each image is presented in the front view side view, and the variable region is zoomed in to highlight the variability. The angle of rotation mentioned is calculated where the α chain is fixed on a conformer, and the angle of rotation is calculated for β and β’ chain.

### 2.5. Conformer E adopts a radical transfer–permissive conformation

**Supplementary Table 4** shows the interface area in the previously reported α_2_β_2_ complexes and the different Mth conformers ^16,38^. The interface area of interaction between α and β is in the order of *E. coli* (Class Ia)> Conformer E> Conformer G> Conformer C> Conformer D> *S. typhimurium*> Conformer B> *B. subtilis* > Conformer A> Conformer F. Among the seven conformers in our structure, the interface area between α and β is maximum in conformer E (2094 Å^2^) whereas the *E. coli* structure with mechanism-based inhibitor N3CDP shows the highest interface area (3634 Å²) ^40^. On the other hand, the conformer F shows minimum area of interaction between α and β i.e. 825 Å².

The distance from the metal cofactor site in β chain (Tyr 106) to the active site (Cys 367) in α chain among the seven conformers varies between 50 Å to 37.6 Å **(Supplementary Figure 5A),** and the same for β chain to α’ chain varies from 63 Å to 41 Å **(Supplementary Figure 5B)**. The variations in overall structure and inter-subunit distances are illustrated for Mth **(Figure 3A and 3E),** E. *coli (*class Ia) **(Figure 3C and 3G),** and the class Ib structures of *S. typhimurium* **(Figure 3B and 3)** and *B. subtilis* **(Figure 3D and 3H).** In conformer E, the β–α and β–α′ distances are 37.6 Å and 42 Å, respectively. Conformer F resembles the active state *E. coli* RNR structure, with an α–β distance of 41.7 Å, compared to 37.6 Å in conformer E, 37.8 Å in *B. subtilis*, and 32.4 Å in *E. coli*. The relatively shorter distance in conformer E suggests that it requires minimal rearrangement and may facilitate radical transfer. Since conformer E has the maximum interface area and the shortest path for radical transfer from the metal cofactor site to the active site, it appears to be the closest to the active state conformation among the seven conformers **(Figure 3A).** Although, these structures lack substrate, this enzyme inherently samples radical transfer–permissive conformation conformations even in the absence of ligand.

**Figure 3:**
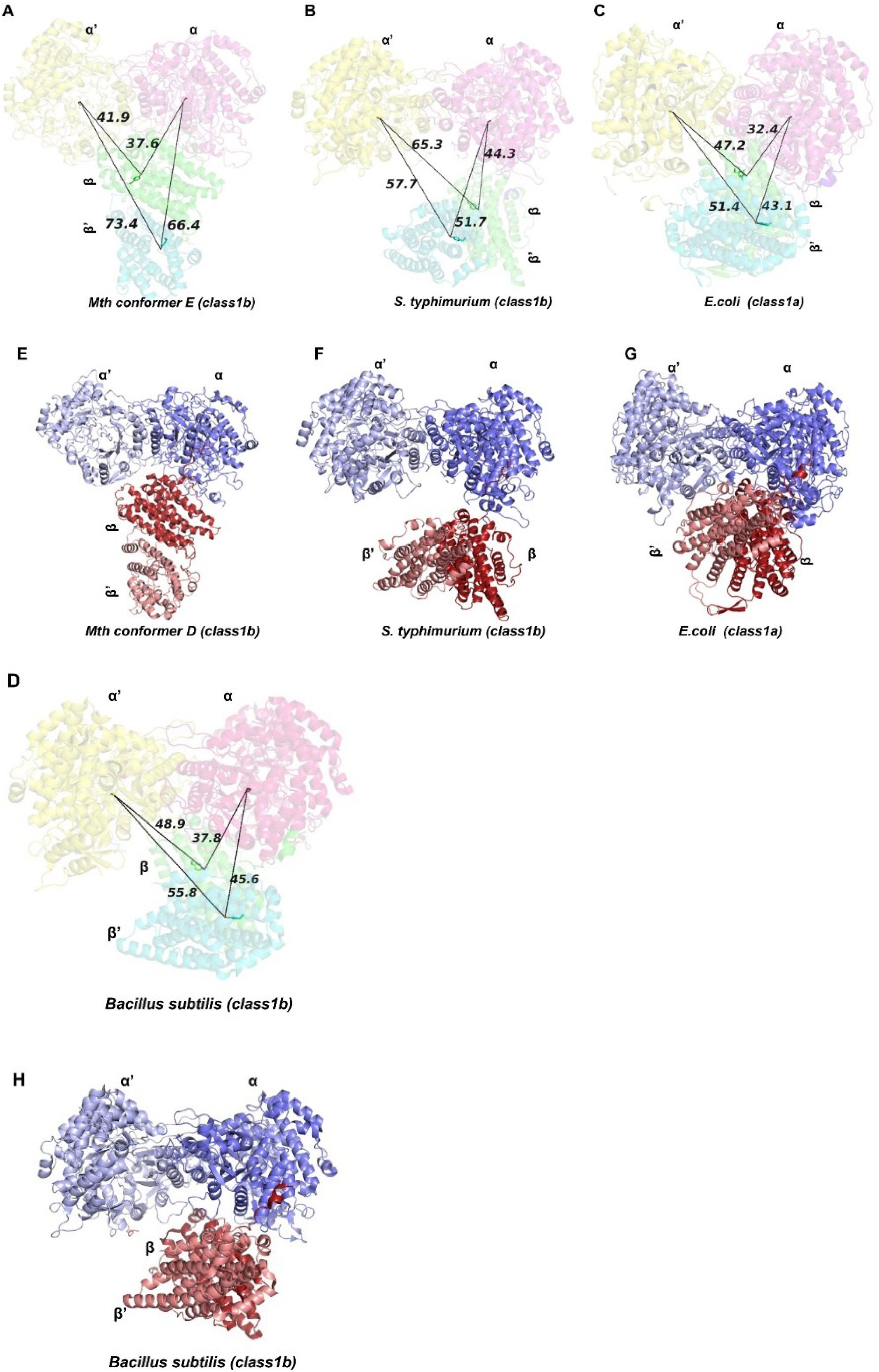
Variability in α_2_β_2_ complexes. Comparative analysis of variability observed in radical transfer distance from α subunit to β subunit in different α_2_β_2_ complexes. The distance from tyrosine at the metal cofactor site to cysteine at the active site is measured from β to α, β’ to α, β’ to α, and β’ to α’ chain for **A.** Conformer E, **B.** *S. typhimurium* (2BQ1), **C.** *E. coli* (6W4X), **D.** *B. subtilis* (9BZ9). The distance is shown in Å for the different conformers as labelled. Cartoon representation of the structure of α_2_β_2_ complex to demonstrate variability in presented structure of **E.** Mth Conformer D class Ib**, F.** *S. typhimurium* class Ib**, G.** *E. coli* class Ia, **H**. *B. subtilis* class Ib.

### 2.6. Near-atomic resolution of conformer D reveals the architecture of the α_2_β_2_ complex

In the 3.8 Å structure of conformer D, we can trace all the four chains with a good model-map correlation, except for some known disordered regions **(Table 1).** However, α’ exhibits a poor map-model correlation at its N-terminal region **(Supplementary Figure 6A).** Density for ∼100 residues was missing at N-terminus, therefore, 105 residues from the N-terminal region of the α’ chain were not modelled. The density for the last 20 residues in the C-terminal is also unresolved and is not present in both protomers of the α subunit. Apart from the two termini and the loop regions, the main chain for most parts of the α and α’ chains are clearly visible, and the side chain density is also resolved in some segments. In the β chain, we were able to trace the density of the entire chain, including the C-terminal region although at this region density was poor **(Supplementary Figure 6B).** On the other hand, the β’ chain has ∼27 residues missing at the C-terminal region. In the β and β′ subunits, low-resolution density is observed at the metal-cofactor site. Previous work using X-ray fluorescence scan of the NrdF2I crystal confirmed the presence of Mn at this site without external metal supplementation, so we modelled two Mn²⁺ atoms at this site ^26^**. Supplementary Figures 6C-6J** show a few representative images demonstrating density details from different chains in the complex.

The asymmetry observed in conformer D is similar to previously reported RNR structures (**Figure 4)** ^16,38,1^. The hinge-like attachment between β and α subunits is evident, indicating significant inter-subunit flexibility **(Figure 4A and 4B).** In conformer D, the distance between the metal cofactor site tyrosine (β chain) and the active site cysteine (α chain) is 43 Å in α chain and 54 Å in α’ chain. **Figure 4C** shows the tyrosine cluster and cysteine at the active site in the α chain, the density for which is clearly visible **(Figure 4D).** Conformer D is the only class Ib α_2_β_2_ complex structure where the low-resolution density of β chain C-terminal region is observed in its native ligand-free state compared to reported structures **(Figure 4E). Figure 4F** illustrates the electron density corresponding to side-chain residues of the C-terminal tail region of the β chain.

**Figure 4:**
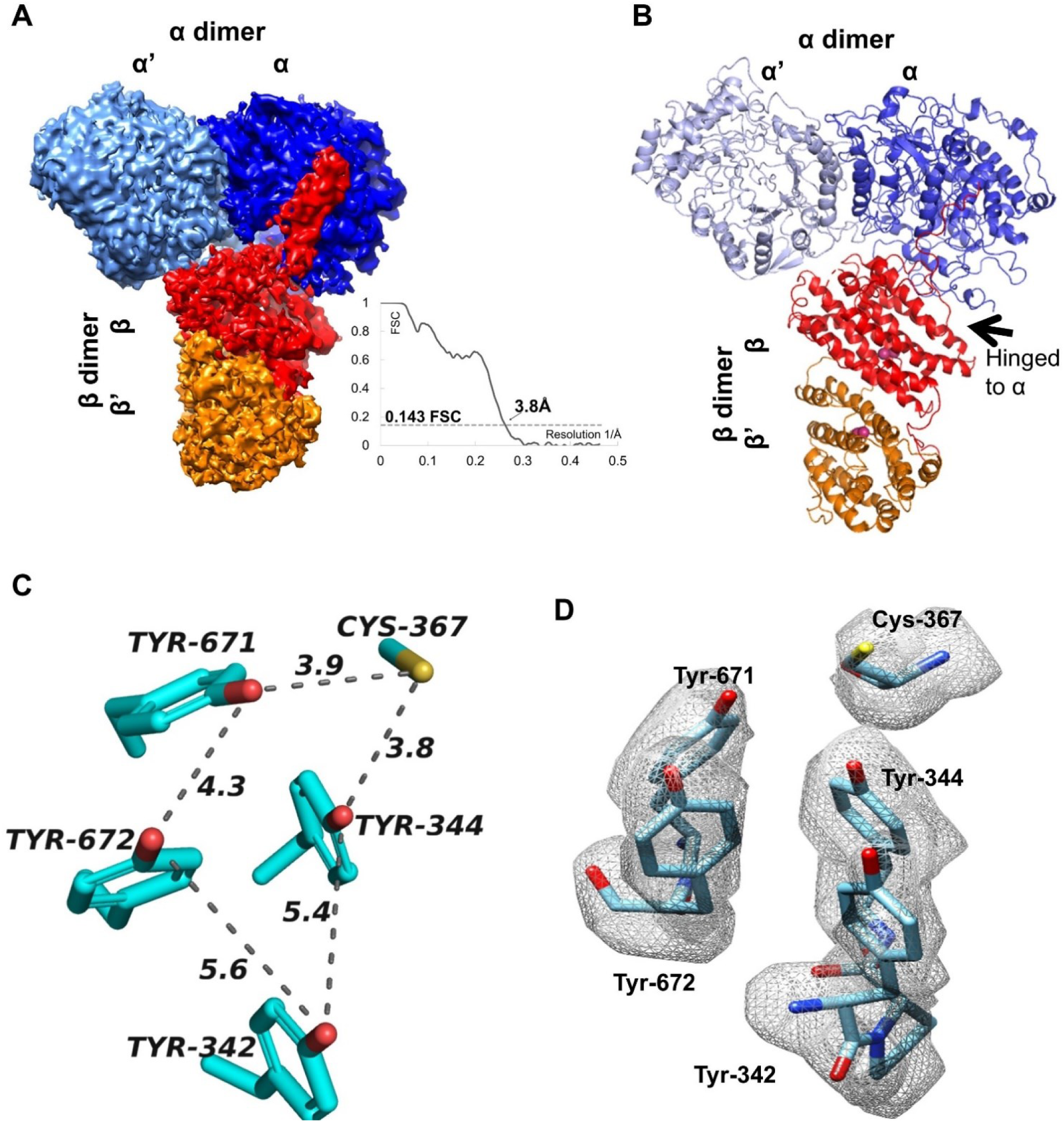
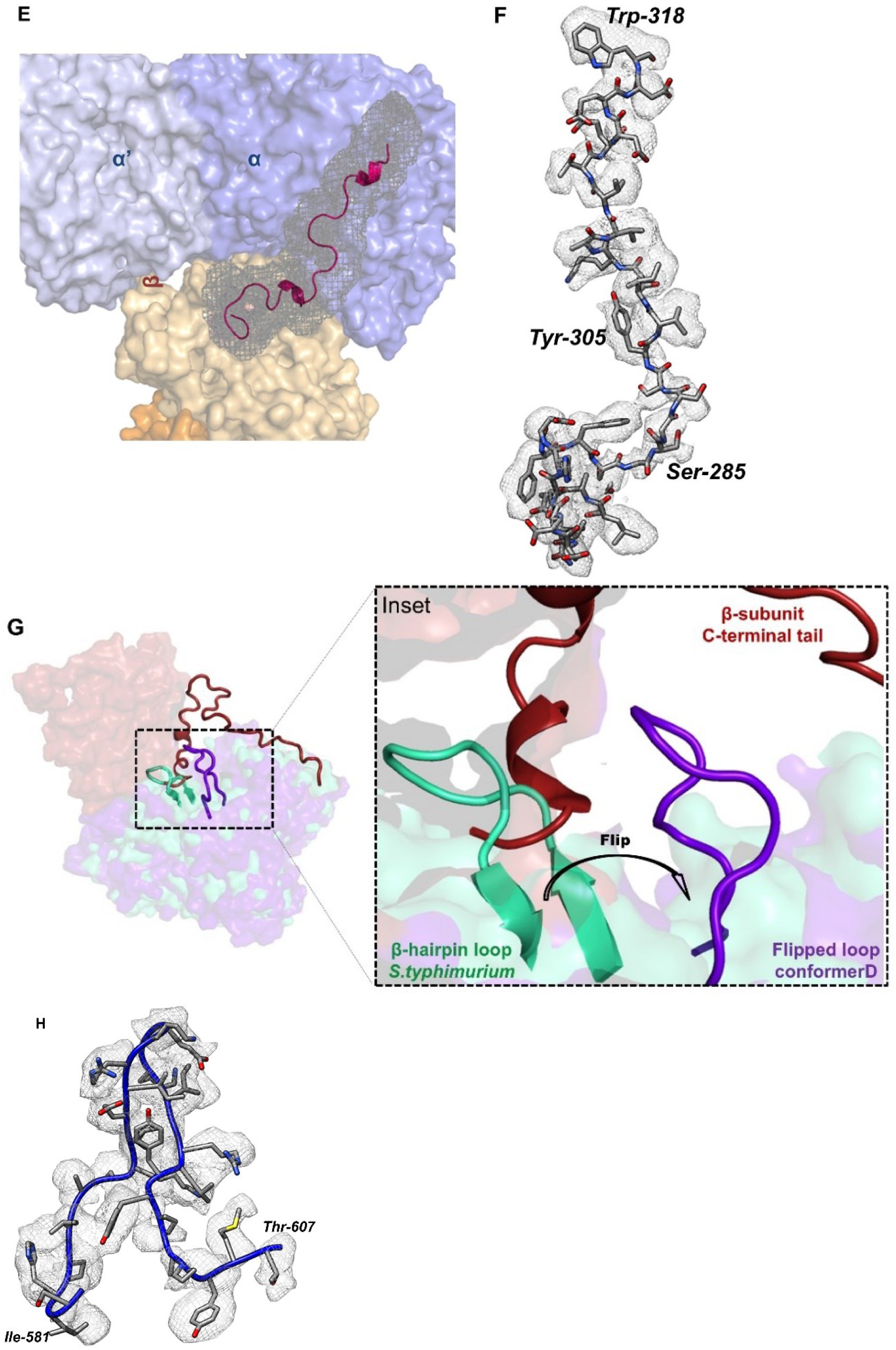
Cryo-EM structure of Conformer D of Mth α_2_β_2_ complex. **A**. CryoSPARC sharpened cryo-EM density map with dark/light blue color represents α subunit, and red /orange represents β subunit. The map was contoured at 0.2σ. Inset shows the FSC curve for conformer D at 0.143 FSC. **B.** Structure of α_2_β_2_ complex. **C.** Residue at the active region in α subunit demonstrating tyrosine cluster and cysteine present at the active site. **D.** Active site cryo-EM density of the residues involved in nucleotide reduction demonstrated using density modified map. This density modified map was generated using Resolve CryoEM in Phenix, using only the half map and the unsharpened full map, with no model provided during the process. The map was contoured at 0.05σ. Colour zone tool of chimera with radius cutoff of 1.5Å was used to map the active site density. **E**. Unsharpened map density (black mesh) highlighting the C terminal tail region of β chain (pink cartoon). **F** CryoSPARC sharpened map demonstrating density for side chain residues of C-terminal tail region of β chain. The map was contoured at 0.1σ. and to map density radius cutoff of 2Å was used. **G.** Surface representation of α and β chain of superposed *S. typhimurium* and conformer D structure. Inset highlighting the β hairpin loop of *S. typhimurium* (green) clash with the C-terminal tail of the β subunit of conformer D (red) and flip observed of a similar loop (blue) in conformer D. **H.** Cryosparc sharpened map (B factor of -155.5 Å^2^) highlighting the density of β hairpin loop with map contoured at 0.1σ. Colour zone tool of chimera with radius cutoff of 2Å was used to map the density.

A remarkable feature of conformer D is the pronounced conformational shift in the β hairpin loop of the α subunit comprising residues 586–600, compared to the equivalent region in *S. typhimurium* **(Figure 4G). Figure 4H** shows the density of this β hairpin loop. This loop undergoes a displacement of approximately 16 Å generating a cavity within the α subunit that accommodates the C-terminal tail of the β subunit **(Figure 4G).** Our structure suggests that this movement not only allows access of the first half of the β C-terminal tail into the binding pocket, but that flipping of the loop further facilitates interactions with the distal portion of the C-terminal tail, effectively locking the β subunit onto the surface of the α subunit **(Figure 4G inset) (Supplementary video 2).** A similar kind of flexibility of β hairpin loop has been reported in the α subunit structure of *E. coli* ^2^, however, the functional role of this loop in presence of β subunit in α_2_β_2_ remained unclear. On the contrary, the β-hairpin loop (642 to 655) from the α subunit in *E. coli* swings towards the active site and stabilises the C-terminal of the β subunit for radical transfer ^16^. Comparable structural transitions are also observed in class II RNRs, where conformational changes are induced in a B12-induced manner ^41^, suggesting a conserved role for loop dynamics in facilitating substrate access and catalysis across RNR classes.

### 2.7. Interactions of C-terminal tail of β subunit in α_2_β_2_ complex

The C-terminal tail of the β chain (residues 270–320) can be divided into two segments: proximal (∼270–307) and distal (∼308–320) segments **(Supplementary Figure 8A).** In six of the seven conformers (B, C, D, E, F, and G), poor density for distal part of the C-terminal tail is present, anchored on the surface of α chain, similar to that observed in *S. typhimurium* and *B.subtilis* structure **(Supplementary Figure 8B).** Density for the proximal part of the C terminal is weak and discontinuous in conformers A, B, C, and E **(Supplementary Figure 8E, 8F, 8G, and 8I),** while conformers F and G show the presence of weak and continuous density **(Supplementary Figure 8J and 8K).** On the other hand, in conformer D, the entire C-terminal tail is seen in the map, where the proximal tail extends into the active site of the α chain, while the distal end attaches on the surface of the α chain **(Supplementary Figure 8A and 8H).** Moreover, the flexibility of the C-terminal region near the active site is evident from the low-resolution/ fragmented density observed in this region in all the conformers. The complete C-terminal tail in β subunit has previously been observed only in the Class Ia cryo-EM structure of *E. coli* RNR ^16, 40^ **(Supplementary Figure 8C).** Our structure therefore provides the first view of the C-terminal tail orientation in the ligand-free β subunit of class Ib RNR despite the limited resolution. Interestingly, the orientation of the C-terminus observed in classes Ia and Ib is distinctly different in the proximal part. The relative orientation of β chain C-terminal tail, when only α-chain **(Supplementary Figure 8M),** or only β-chain (**Supplementary Figure 8L**) is superposed from the α_2_β_2_ complexes of *E. coli* and Mth, is shown. Both superpositions revealed notable variations in the C-terminal tail, consistent with differences in the orientation of the β-subunit between the two complexes.

Three-dimensional variability analysis (3DVA) of conformer D revealed heterogeneity consistent with changes in the position of the β C-terminal region relative to the active site of the α chain ^42^. To evaluate the extent of movement, low-resolution model building was performed. Model building of two extreme maps of components 0 and 1 of 3DVA indicate dynamic reorientation in the C-terminal region away from the active site **(Figure 5A and D)** or towards the active site of the α chain **(Figure 5B and E**). **Figure 5C and F** depict a map where both structures are superimposed, illustrating the observed differences in the tail region. Locating the residues in this C-terminal region indicates that Tyrosine 305 of the β chain can adopt orientation closer to the tyrosine cluster near the active site in the α chain **(Figure 5G).** A corresponding tyrosine residue in class Ia is considered to be important in radical transfer and is the last residue of the β subunit in radical transfer steps ^18,17^. This observation is therefore consistent with, but does not directly demonstrate, a potential role in radical transfer

**Figure 5:**
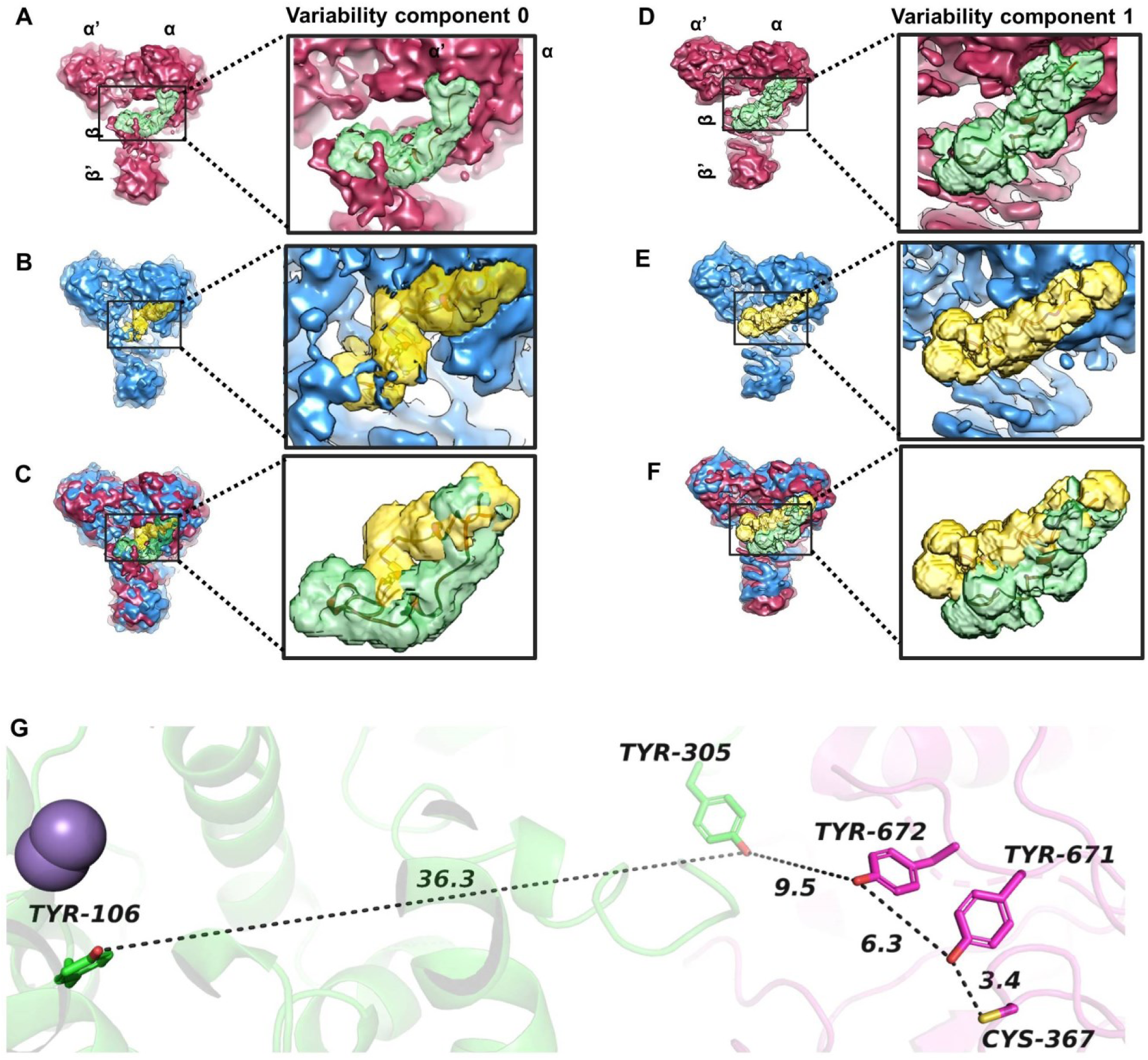
Evaluation of C terminal tail flexibility of β chain using 3DVA map. **A and B.** 3DVA component 0 with two extreme maps demonstrating variation in C-terminal tail in β chain of α_2_β_2_ complex and **C.** shows superposition of maps A and B. **D and E.** 3DVA component 1 with two extreme maps demonstrating variation in the C-terminal tail in the β chain of α_2_β_2_ complex and **F.** shows a superposition of map D and E. C-terminal tail is highlighted in yellow and green. **G.** The model built from this low-resolution 3DVA map indicated movement of β chain C-terminal tail towards α chain, decreasing the distance of tail tyrosine 305 from tyrosine cluster (aa 671 and 672) at the active site by ∼10Å. Extreme maps are the positive and negative standard deviation projections along the principal component of variance for component 0 and 1.

In addition to C-terminal flexibility, the influence of global β-subunit motion on the α–β distance in the α₂β₂ complex was evaluated. The distance between α Cys367 and β Tyr106 varies from 50.5 to 37.6 Å **(Supplementary Figure 5A),** with conformers A/B showing ∼50 Å and conformer E showing 37.6 Å. This distance may be further reduced by C-terminal tail movement of the β subunit, as observed in 3D variability analysis of conformer D **(Figure 5)**.

### 2.8. The α_2_β_2_ interaction is enthalpically-driven

To investigate the thermodynamics of α and β subunit interaction, we conducted a qualitative binding analysis of the α subunit to immobilized β subunit at different temperatures using SPR **(Supplementary Figure 9A).** The overlay of sensogram at different temperatures indicated that the binding kinetics slowed with a decrease in temperature, suggesting a temperature-dependent reaction **(Supplementary Figure 9B).** The dissociation and association kinetics of α and β subunits were estimated at different temperatures of 10-30°C at 4°C intervals. The equilibrium dissociation constant, determined for the α_2_β_2_ complex, varied from 9.6 × 10^−8^ M (10°C) to 4.1 × 10^−7^ M (30°C). An increase in temperature was observed to decrease the affinity of α to the immobilised β subunit. Kinetic analysis revealed the decrease in affinity observed is primarily due to a reduction of ka with a slight increase in kd

The thermodynamic parameters estimated provide insights into the overall changes occurring during the interaction. The bar graph indicates that the reaction is exothermic and accompanied by a decrease in entropy upon complex formation **(Supplementary Figure 9C).** The high entropic cost observed in SPR likely reflects the disorder-to-order transition of the β-tail observed in Conformer D. **Supplementary Figure 9D** shows the plot of Gibbs free energy change as a function of temperature. This change in ΔG with temperature suggests that β and α subunits may undergo significant conformational changes upon binding. Specifically, the binding site in the protein might experience a conformational adjustment during the interaction. To conclude, the interaction of α and β subunits is an enthalpically driven interaction brought about by stabilizing a flexible region combined with markedly unfavourable entropic forces.

## 3. Discussion

RNR complexes have historically been difficult to structurally characterize due to their intrinsic dynamics and weak, transient subunit interactions. Although several efforts using inhibitors and unnatural amino acid incorporation have been made to stabilize and trap the holoenzyme, a complete structural description has remained challenging^43,29,16^. Here, we present the wild-type ligand-free structure of the *M. thermoresistibile* (Mth) α₂β₂ complex belonging to the class Ib RNR. In *E. coli* (class Ia) RNR, the transient α₂β₂ interaction is characterized by a Kd of ∼400 nM ^44^, while SPR studies in *B. anthracis* (class Ib) report an affinity of ∼178 nM ^45^. In contrast, the Mth α₂β₂ complex exhibits a higher affinity of ∼85 nM at ∼25 °C, resulting in a more stable complex that enabled co-purification and structural characterization (Supplementary Figure 9). Previous stabilization strategies, including β2 double mutants, substrate/effector combinations and inhibitors, have been used to reduce flexibility and trap the holoenzyme ^43,29,16^ but a native ligand-free structure has remained elusive. In contrast, here we directly purified the native complex without stabilizing modifications or reconstitution. Using cryo-EM, we not only obtained a homogeneous ligand-free complex but also determined its structure in multiple conformational states for the first time. This provides a direct structural view of the apo assembly and reveals its intrinsic conformational variability.

### 3.1. Conformational diversity in α_2_β_2_ complex

The 3.8 Å cryo-EM structure reveals that the α and β subunits form an asymmetric heterodimer, consistent with previous X-ray and cryo-EM structures ^16,38^. **Table 2** provides an overview of previously reported α₂β₂ complex RNR structures. Interestingly, none of the α₂β₂ complexes from different organisms align with one another, exhibiting significantly different modes of association between the α and β subunits. The Class Ib α₂β₂ complexes from *S. typhimurium* and *B. subtilis*, both determined in the presence of nucleotides, also show markedly different modes of subunit association, with the interface exposing a larger surface area to the external solvent ^38^ (**Figure 3 and Supplementary Table 4**). These differences likely arise from the intrinsically dynamic and transient nature of the α-β interaction, and ligand-dependent conformational changes, which suggests that different structures capture distinct functional or conformational states of the complex. Moreover, our data also reveals seven distinct conformational states highlighting local and global flexibility and the dynamic disorder within the α_2_β_2_ complex. The unprecedented extent of variability between the two subunits observed in our complex has not been seen earlier in any of the reported structures, including those of different dATP inhibited and cryo-EM structures of class Ia RNR ^16,46,47^. Recently, Xu et al. applied cryo-EM combined with deep-learning-based methods to the Class Ib α₂β₂ complex from *B. subtilis*, revealing that the enzyme samples continuum of conformations. It samples wide range of motions from open to compact including PCET-competent conformation observed in *E. coli* Class Ia RNR ^39^. SAXS and structural investigations of the class Ib α_2_β_2_ complex in *Bacillus subtilis* reveal a range of conformational ensembles in solution in different conditions, including a flexible tetramer and inhibited filament structures^1^.

**Table 2:**
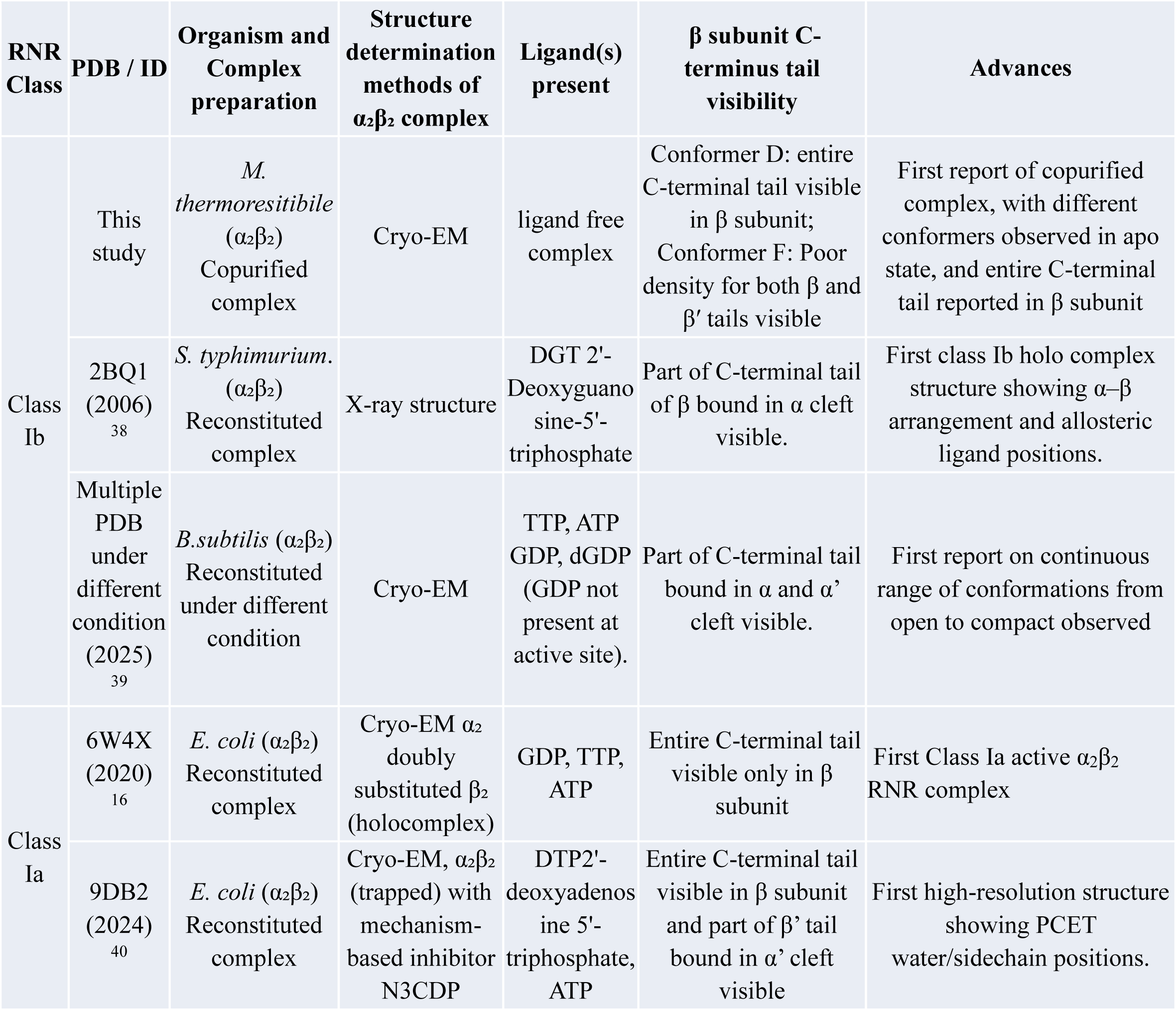
Previously determined X-ray and Cryo-EM structures of α_2_β_2_ complex of Class Ia and Class Ib RNR.

We collected cryo-EM data for the ternary complex NrdEFI multiple times and consistently observed similar conformational variability, indicating that this heterogeneity reflects intrinsic dynamics of the complex rather than experimental artefacts. However, particle heterogeneity limited the achievable resolution. To mitigate this, we acquired data on crosslinked binary (α₂β₂) complex. Notably, this approach yielded an enriched dominant conformational state, resulting in improved map quality and higher resolution. Despite the absence of NrdI in binary α₂β₂ complex, it retained conformational variability similar to that observed in the ternary complex (NrdEFI), indicating that much of the flexibility is inherent to the NrdEF or α₂β₂ core. Among the seven conformers observed, conformers A and B with less interface area between α and β subunits adopt an extended conformation and might represent a transient initial stage of complex formation. Conformer E adopts a compact conformation with maximum interaction interface area **(Supplementary Table 4),** while conformers C, D, F, and G adopt intermediate conformations. In conformer F, the β chain associates with the α chain in a slightly tilted manner compared to other intermediate conformations. Although not identical, conformer F most closely resembles the structures of *E. coli (*class Ia), *S. typhimurium and B.subtilis (*class Ib) RNRs ^16,3839^. Interestingly, Conformer F is the only conformer where poor density for C-terminal in both the β and β’ chain is observed.

Multiple proteins, such as ribosome^48^, ATP synthase^49,50^ etc. exist in multiple states which correspond to active, inactive, slow and fast activity states. Similarly, proteins involved in DNA replication and repair show distinct conformations that regulate activity, substrate recognition, and so on thereby ensuring speed, fidelity, and genome stability ^51,52^ ^53,54^ ^55^. During DNA replication, billions of deoxynucleotides are required (humans ∼6 × 10⁹; *E. coli* ∼9.2 × 10⁶ nucleotides), whereas DNA repair typically involves the incorporation of only a few hundred nucleotides ^56^. These large differences in deoxynucleotide demand suggest that distinct modes of α₂β₂ subunit association observed may correspond to different functional states of the enzyme. Conformations resembling the *E. coli* complex are likely to represent a high-activity state optimized for rapid nucleotide production during replication. In contrast, structures similar to conformer E may correspond to a low-activity state, potentially suited for conditions such as DNA repair, where only limited nucleotide incorporation is required. Based on these functional distinctions we may speculate that structural variability among RNR complexes reflect adaptation to the cellular demand for deoxyribonucleotide synthesis.

### 3.2. Relative ordering of the proximal segment of the β subunit C-terminal tail

In the α₂β₂ complex, the primary interaction interface involves the α N-terminal domain and the β C-terminal (∼50 residues), which are often poorly resolved or absent in isolated subunits. Complex formation stabilizes the α N-terminus, as observed upon substrate binding in holo Mth α dimer structure ^37^. In contrast, the α′ N-terminus remains poorly defined in both X-ray and cryo-EM structures ^16, 37^. The absence of clear density in the α′ chain therefore likely reflects increased flexibility, which may provide a functional advantage by enabling conformational sampling. Similarly, the β C-terminal tail is frequently unresolved in prior X-ray and NMR studies ^216,38,57^. In α₂β₂ structures from *B. subtilis* and *S. typhimurium*, only the distal portion of this tail is visible, whereas the full C-terminal tail is resolved in the *E. coli* class Ia cryo-EM structure ^16^. This structure positions Tyr356 for proton-coupled electron transfer (PCET) and implicates the tail in substrate docking and product release ^16^.

Conformer D is the only class Ib structure in which the entire β C-terminal tail is resolved. The distal segment anchors to the α subunit, stabilizing the complex, whereas the proximal half shows relative ordering so that it is critically placed closer to the active site in the α chain **(Figure 5).** This is mediated by an α-subunit β-hairpin loop that in *S. typhimurium* occupies the same pocket as the proximal β tail, creating a steric clash, whereas in conformer D it adopts a flipped-out conformation that resolves this conflict **(Figure 4G and Supplementary video 2)**. This β-hairpin loop has been previously implicated in substrate stabilization and is known to adopt distinct conformations in the substrate-bound and substrate-free states ^2^. We propose that substrate alignment triggers β-hairpin loop repositioning, enabling engagement of the proximal half of the β tail and supporting a gating-like rearrangement consistent with PCET regulation. Thus, the putative mechanism of proton-coupled electron transfer is revealed in an unprecedented detail.

### 3.3. Mechanistic insights into β’ and α’ interaction

The dynamic nature of the α₂β₂ complex association provides insight into the interaction between the α′ and β′ chains of the heterotetramer. The movement of the β subunit relative to the α subunit observed across conformers suggests a mechanism that not only modulates α–β interactions but may also facilitate reorganization of the α′ and β′ chains. This interaction is relevant for subsequent catalytic cycles, where β′, carrying the radical, would transfer it to α′. Thus, the observed dynamics suggest a possible reorientation leading to α′–β′ proximity.

The distinct conformers obtained from our cryo-EM data represent snapshots of transient states of the α₂β₂ complex. These can be grouped into structural regimes consistent with stages of the catalytic cycle, although their temporal ordering remains hypothetical: (1) rotational movement of the β subunit towards the two-fold symmetry axis of the α subunit (conformers A–D and G); (2) compaction of the β subunit towards α (conformer E); and (3) forward movement of the β′ chain towards α′ (conformer F) **(Figure 6). Supplementary Video 4-7** depicts our hypothesis on the proposed major transitions observed in different maps through morphing.

**Figure 6:**
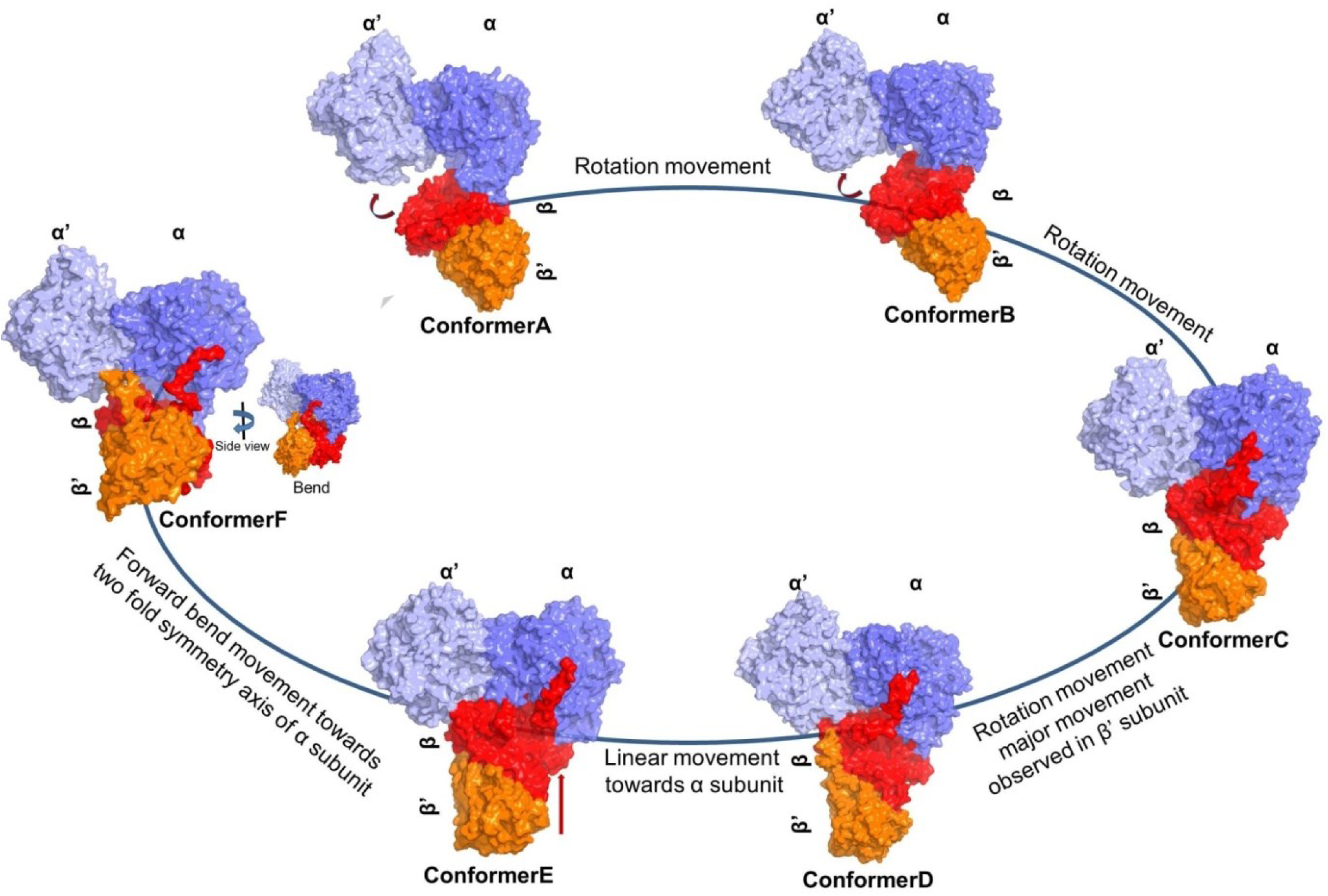
Proposed mechanism illustrating conformational transitions in the α₂β₂ complex and probable pathways leading to α′ and β′ chain association. The different conformers are arranged based on available structural information on RNR and the movement of the β subunit relative to the α subunit. These conformers suggest distinct modes of β–α interaction, providing a structural framework for the proposed mechanism of protein function and a potential pathway for α′ and β′ chain association after reduction. Blue/light blue denotes α/α′, and red/orange denotes β/β′ subunits.

In conformers A and B, the substrate-binding site remains accessible, although these open states are incompatible with radical transfer. However, in these conformers their active sites remain accessible to the C-terminus of the α chain, enabling reactivation of active site disulphide. In conformers A–D, rotation of the β subunit towards α is observed and may be coupled to substrate-induced stabilization of the α N-terminus ^37^**(Figure 6).** Concurrently, the α-subunit β-hairpin loop shifts away from the active site to accommodate the β C-terminus. In conformer E, compaction of the β subunit and repositioning of its C-terminal tail towards the α active site may facilitate radical transfer and cysteinyl radical formation **(Figure 5).** A similar mechanism has been proposed for class Ia RNR, where the β-tail regulates substrate access during proton-coupled electron transfer. Upon product formation, disulfide formation in α induces strain that displaces the β-tail, enabling product release and exposing the disulfide for re-reduction, thereby resetting the enzyme for the next catalytic cycle^16^. Subsequently, coordinated movement of β and β′ (conformer F) brings the α′ and β′ chains into closer proximity while weakening α–β interactions, potentially facilitating product release and enabling formation of an alternative interface. A transient state following conformer F may resemble the *E. coli* class Ia structure, where a closer α′–β′ interaction is observed ^16^ **(Figure 6 and Supplementary video 8).** This proposed sequence is consistent with known steps in ribonucleotide reduction. **Supplementary Video 9** illustrates a hypothetical gradual transition between conformers, highlighting the interaction mechanism between α and β. The video also highlights the probable mode of association between α’ and β’, ultimately culminating in a structural state resembling the *E. coli structure*. This proposed sequence is consistent with the known steps for reduction of ribonucleosides. Our study of the ligand-free Mth complex defines the baseline conformational ensemble accessible to the enzyme. The observed conformational heterogeneity may reflect kinetic memory, as proposed for persistent α₂β₂ association in *E. coli* and the α₆ state in human RNR ^43^. However, without time-resolved or kinetic data (e.g., MD simulations, FRET, HDX-MS), the ordering remains hypothetical, and some states may represent low-population or off-pathway intermediates.

To conclude, we have obtained important insights into the multiple conformational states of native Mth RNR complex through cryo-EM single particle analysis. The structural flexibility and conformational variability observed in the α_2_β_2_ complex are essential for reducing ribonucleoside to deoxyribonucleoside diphosphates, a critical step in DNA synthesis. This study enhances our understanding of the ribonucleotide reductase and highlights the importance of structural dynamics in enzyme function.

## 4. Materials and methods

### 4.1. Purification of Mth α_2_β_2_ complex

The α_2_β_2_ /(NrdEF) binary and NrdEFI ternary complexes were co-expressed and co-purified. The soluble fraction of α_2_β_2_ complex was purified using affinity and size-exclusion chromatography. Details of purification and crosslinking of α_2_β_2_ are provided in **Supplementary Section 1.2 and 1.3.** The thermodynamic parameter of α and β interaction at different temperatures using SPR is described in the **Supplementary section 1.6.**

Briefly, NrdEFI expressing cells were lysed in a Tris-based buffer containing 300 mM NaCl, 3% glycerol, 0.5 mM EDTA, 1 mM MnCl₂, and protease inhibitors. The complex was then purified from clarified lysate using Ni-NTA affinity chromatography. On column cleavage with thrombin was used to remove tag from the α and β subunits. The resulting NrdEFmbpI-ternary complex was concentrated and further purified by tandem size-exclusion chromatography on 16/600 Superdex 200 and 10/300 Superdex 200 increase column in buffer Tris 25 mM pH 7.5, NaCl 150 mM, 1mM MnCl_2_.

### 4.2. Grid preparation and data acquisition

Purified NrdEFI (2 mg/ml) and cross-linked α_2_β_2_ (1.25 mg/ml) complexes were applied (3 µl) to glow-discharged Quantifoil Au R0.6/1 (300 mesh) grids, blotted, plunge-frozen, and stored in liquid nitrogen. Cryo-EM data for the ternary complex were collected at ESRF (CM01) on a Titan Krios G3 (300 kV, K2 detector) at 130K magnification (1.052 Å pixel size), yielding conformers A, C, E, F, and G. Data for the α_2_β_2_ complex were acquired at the National Cryo-EM facility, Bangalore, on a Titan Krios G3i (300 kV, K2 detector, energy filter) at 130K (1.065 Å pixel size), yielding conformers B and D. The details of grid preparation and data acquisition is described in detail in **Supplementary section 1.4.**

### 4.3. Cryo-EM image processing and classification

Beam-induced motion was corrected using MotionCor2 in RELION, and subsequent processing was carried out in CryoSPARC v3.3.2. Micrographs were CTF-estimated, curated (>10 Å), and particles were template-picked, filtered, and subjected to multiple rounds of 2D classification to remove bad particles. Selected particles were re-extracted, duplicates removed, and used for ab initio reconstruction, heterogeneous refinement, and 3D variability analysis. Ab initio classification provided better separation of conformations, leading to 12 classes for the ternary complex, while the binary complex was refined using selected volumes. Final high-resolution maps were obtained via non-uniform and local refinement without imposed symmetry, with resolution and map quality assessed using FSC, angular distribution, and local resolution estimation. The details are discussed in **Supplementary section 1.5**

For Conformer D (3.8 Å), a cryoSPARC map sharpened at a B value of -155.5 Å^2^ was used. Autosharpened maps of conformers A, B, C, E, and G, DeepEMhancer ^58^ map for conformer F were used for model docking and interpretation of global conformational change. Unsharpened maps were used to demonstrate the path of the flexible C-terminal tail of the β subunit. For analysis, the data that was good for the interpretability of the map was used for model building, as the division of particles into different classes compromised the resolution of the map **(Supplementary Figure 1).**

### 4.4. Model building and refinement

To determine the structure of the complex, Cryo-EM maps were docked with models obtained from the crystal structures. The chainsaw model made from the α subunit (PDB ID 2BQ1) and the β subunit (PDB ID 1UZR) was fit through local optimisation in Chimera using the ‘fit in map’ option. Flexible refinement was applied to all conformers following rigid-body docking, but due to limited map quality, detailed model building and refinement were performed only for conformer D used in the final model. These individual models were combined to get a binary complex followed by flexible fitting in iMODFIT ^59^ and Namdinator ^60^. The model thus obtained was further refined in Phenix with real space refine with secondary structure and geometry restraints. Multiple rounds of refinement in Namdinator and Phenix (v1.20.1) were done to improve the fitting of the map and model. Model-map FSC for conformer D was estimated to be 4.4 Å at a threshold of 0.5. The geometry of the final model was evaluated using MolProbity ^61^. All figures and images were made in Pymol ^62^ and Chimera ^63^.

### 4.5. PDB ID numbers and deposited maps of the structure determined

The coordinates of the Mth α_2_β_2_ complex (conformer D) are deposited in a protein data bank under PDB ID - 9L6O with accession codes EMD-62860 of the associated cryo-EM maps. The Cryo-EM maps of Conformer A, Conformer B, Conformer C, Conformer E, Conformer F, and Conformer G are deposited in EMDB with EMD nos EMD-62854, EMD-62858, EMD-62855, EMD-62857, EMD-62859 and EMD-62856 respectively.

## Acknowledgments

This work was supported by a DBT-Centre of Excellence Grant (BT/PR15450/COE/34/46/2016) and DST-NPDF (PDF/2016/000774). Genomic DNA of *M. thermoresistibile* was provided by Dr. Christoph Grunder (University of Washington). Cryo-EM data were collected at the inStem Bangalore facility (DBT/PR12422/MED/31/287/2014), with thanks to Dr. Vinothkumar Kutti for access, grid preparation and discussions. Synchrotron beam time was provided by ESRF (CM01), with assistance from Daouda Traore. SPR experiments were performed at NCCS Pune, with support from Venkatesh Rajmane and Tejashree Damale. PT acknowledges DBT Research Associateship and DST NPDF. SCM and LRY acknowledge DST JC Bose Fellowship; SCM also acknowledges support from Anand Deshpande (Savitribai Phule Pune University).

## References

1. Thomas, W. C. et al. Convergent allostery in ribonucleotide reductase. Nat. Commun. 10, (2019).

2. Zimanyi, C. M., Chen, P. Y. T., Kang, G., Funk, M. A. & Drennan, C. L. Molecular basis for allosteric specificity regulation in class ia ribonucleotide reductase from Escherichia coli. Elife 5, 1–23 (2016).

3. Minnihan, E. C., Nocera, D. G. & Stubbe, J. A. Reversible, long-range radical transfer in E. coli class Ia ribonucleotide reductase. Acc. Chem. Res. 46, 2524–2535 (2013).

4. Reece, S. Y. & Seyedsayamdost, M. R. Long-range proton-coupled electron transfer in the Escherichia coli class Ia ribonucleotide reductase. Essays Biochem. 61, 281–292 (2017).

5. Nordlund, P. & Reichard, P. Ribonucleotide reductases. Annu. Rev. Biochem. 75, 681–706 (2006).

6. Ruskoski, T. B. & Boal, A. K. The periodic table of ribonucleotide reductases. Journal of Biological Chemistry 297, 101137 (2021).

7. Torrents, E. Ribonucleotide reductases: Essential enzymes for bacterial life. Front. Cell. Infect. Microbiol. 4, 1–9 (2014).

8. Mowa, M. B., Warner, D. F., Kaplan, G., Kana, B. D. & Mizrahi, V. Function and regulation of class I Ribonucleotide reductase-encoding genes in Mycobacterial. J. Bacteriol. 191, 985–995 (2009).

9. Andersson, C. S. & Högbom, M. A Mycobacterium tuberculosis ligand-binding Mn/Fe protein reveals a new cofactor in a remodeled R2-protein scaffold. Proc. Natl. Acad. Sci. U. S. A. 106, 5633–5638 (2009).

10. Dawes, S. S. et al. Ribonucleotide Reduction in Mycobacterium tuberculosis: Function and Expression of Genes Encoding Class Ib and Class II Ribonucleotide Reductases. Infect. Immun. 71, 6124–6131 (2003).

11. Yang, F. et al. Characterization of two genes encoding the Mycobacterium tuberculosis ribonucleotide reductase small subunit. J. Bacteriol. 179, 6408–6415 (1997).

12. Arora, A. et al. Structural biology of Mycobacterium tuberculosis proteins: the Indian efforts. Tuberculosis (Edinb). 91, 456–468 (2011).

13. Crona, M. et al. A ribonucleotide reductase inhibitor with deoxyribonucleoside-reversible cytotoxicity. Mol. Oncol. 10, 1375–1386 (2016).

14. Hamann, C. S. et al. Chimeric small subunit inhibitors of mammalian ribonucleotide reductase: A dual function for the R2 C-terminus? Protein Eng. 11, 219–224 (1998).

15. Greene, B. L. et al. Ribonucleotide Reductases (RNRs): Structure, chemistry, and metabolism suggest new therapeutic targets. 45–75 (2020)

16. Kang, G., Taguchi, A. T., Stubbe, J. A. & Drennan, C. L. Structure of a trapped radical transfer pathway within a ribonucleotide reductase holocomplex. Science (1979). 368, 424–427 (2020).

17. Seyedsayamdost, M. R. & Stubbe, J. A. Site-specific replacement of Y356 with 3,4-dihydroxyphenylalanine in the β2 subunit of *E. coli* ribonucleotide reductase. J. Am. Chem. Soc. 128, 2522–2523 (2006).

18. Seyedsayamdost, M. R., Yee, C. S., Reece, S. Y., Nocera, D. G. & Stubbe, J. A. pH rate profiles of FnY356-R2s (n = 2, 3, 4) in *Escherichia coli* ribonucleotide reductase: Evidence that Y356 is a redox-active amino acid along the radical propagation pathway. J. Am. Chem. Soc. 128, 1562–1568 (2006).

19. Seyedsayamdost, M. R., Yee, C. S. & Stubbe, J. Use of 2,3,5-F3Y-α2 and 3-NH2Y-α2 to study proton-coupled electron transfer in *Escherichia coli* ribonucleotide reductase. Biochemistry 50, 1403–1411 (2011).

20. Johansson, R. et al. Structural Mechanism of Allosteric Activity Regulation in a Ribonucleotide Reductase with Double ATP Cones. Structure 24, 906–917 (2016).

21. Ando, N. et al. Structural interconversions modulate activity of Escherichia coli ribonucleotide reductase. Proc. Natl. Acad. Sci. U. S. A. 108, 21046–21051 (2011).

22. Joseph A. Cotruvo Jr and JoAnne Stubbe. An active dimanganese(III)-tyrosyl radical cofactor in Escherichia coli class Ib ribonucleotide reductase†. Biochemistry 23, 1–7 (2008).

23. Boal, A. K., Stubbe, J. & Rosenzweig, A. C. Structural basis for Activation of Class Ib Ribonucleotide Reductase. Science (1979). 329, 1526–1530 (2010).

24. Roca, I., Torrents, E., Sahlin, M., Gibert, I. & Sjöberg, B. M. NrdI essentiality for class Ib ribonucleotide reduction in Streptococcus pyogenes. J. Bacteriol. 190, 4849–4858 (2008).

25. Hammerstad, M., Hersleth, H. P., Tomter, A. B., Røhr, Å. K. & Andersson, K. K. Crystal structure of Bacillus cereus class Ib ribonucleotide reductase di-iron NrdF in complex with NrdI. ACS Chem. Biol. 9, 526–537 (2014).

26. Yadav, L. R., Sharma, V., Shanmugam, M. & Mande, S. C. Structural insights into the initiation of free radical formation in the Class Ib ribonucleotide reductases in Mycobacteria. Curr. Res. Struct. Biol. 100157 (2024)

27. Seyedsayamdost, M. R., Chan, C. T. Y., Mugnaini, V., Stubbe, J. & Bennati, M. PELDOR spectroscopy with DOPA-β2 and NH2Y-α2s: Distance measurements between residues involved in the radical propagation pathway of E. coli ribonucleotide reductase. J. Am. Chem. Soc. 129, 15748–15749 (2007).

28. Argirević, T., Riplinger, C., Stubbe, J. A., Neese, F. & Bennati, M. ENDOR spectroscopy and DFT calculations: Evidence for the hydrogen-bond network within α2 in the PCET of E. coli ribonucleotide reductase. J. Am. Chem. Soc. 134, 17661–17670 (2012).

29. Climent, I., Sjöberg, B. M. & Huang, C. Y. Site-Directed Mutagenesis and Deletion of the Carboxyl Terminus of Escherichia coli Ribonucleotide Reductase Protein R2. Effects on Catalytic Activity and Subunit Interaction. Biochemistry 31, 4801–4807 (1992).

30. Yee, C. S., Seyedsayamdost, M. R., Chang, M. C. Y., Nocera, D. G. & Stubbe, J. A. Generation of the R2 Subunit of Ribonucleotide Reductase by Intein Chemistry: Insertion of 3-Nitrotyrosine at Residue 356 as a Probe of the Radical Initiation Process. Biochemistry 42, 14541–14552 (2003).

31. Greene, B. L., Taguchi, A. T., Stubbe, J. & Nocera, D. G. Conformationally Dynamic Radical Transfer within Ribonucleotide Reductase. J. Am. Chem. Soc. 139, 16657–16665 (2017).

32. Seyedsayamdost, M. R., Xie, J., Chan, C. T. Y., Schultz, P. G. & Stubbe, J. Site-specific insertion of 3-aminotyrosine into subunit α2 of E. coli ribonucleotide reductase: Direct evidence for involvement of Y730 and Y731 in radical propagation. J. Am. Chem. Soc. 129, 15060–15071 (2007).

33. Ekberg, M., Sahlin, M., Eriksson, M. & Sjö, B.-M. Two Conserved Tyrosine Residues in Protein R1 Participate in an Intermolecular Electron Transfer in Ribonucleotide Reductase*. J. Biol. Chem. 271, 20655–20659 (1996).

34. Sjöberg, B. M., Karlsson, M. & Jörnvall, H. Half-site reactivity of the tyrosyl radical of ribonucleotide reductase from Escherichia coli. Journal of Biological Chemistry 262, 9736–9743 (1987).

35. Erickson, H. K. Kinetics in the Pre-Steady State of the Formation of Cystines in Ribonucleoside Diphosphate Reductase: Evidence for an Asymmetric Complex†. Biochemistry 40, 9631–9637 (2001).

36. Ravichandran, K. R., Minnihan, E. C., Wei, Y., Nocera, D. G. & Stubbe, J. A. Reverse Electron Transfer Completes the Catalytic Cycle in a 2,3,5-Trifluorotyrosine-Substituted Ribonucleotide Reductase. J. Am. Chem. Soc. 137, 14387–14395 (2015).

37. Yadav, L. R., Chauhan, S. B., Joshi, M. & Mande, S. C. Structural basis of half-site reactivity in Class Ib ribonucleotide reductases. bioRxiv 2025.12.21.695763 (2025)

38. Uppsten, M., Färnegårdh, M., Domkin, V. & Uhlin, U. The First Holocomplex Structure of Ribonucleotide Reductase Gives New Insight into its Mechanism of Action. J. Mol. Biol. 359, 365–377 (2006).

39. Xu, D., Thomas, W. C., Burnim, A. A. & Ando, N. Conformational landscapes of a class I ribonucleotide reductase complex during turnover reveal intrinsic dynamics and asymmetry. Nature Communications 2025 16:1 16, 1–14 (2025).

40. Westmoreland, D. E. et al. 2.6-Å resolution cryo-EM structure of a class Ia ribonucleotide reductase trapped with mechanism-based inhibitor N3CDP. Proc. Natl. Acad. Sci. U. S. A. 121, e2417157121 (2024).

41. Sintchak, M. D., Arjara, G., Kellogg, B. A., Stubbe, J. A. & Drennan, C. L. The crystal structure of class II ribonucleotide reductase reveals how an allosterically regulated monomer mimics a dimer. Nat. Struct. Biol. 9, 293–300 (2002).

42. Punjani, A. & Fleet, D. J. 3D variability analysis: Resolving continuous flexibility and discrete heterogeneity from single particle cryo-EM. J. Struct. Biol. 213, 107702 (2021).

43. Hassan, A. Q., Wang, Y., Plate, L. & Stubbe, J. Methodology To Probe Subunit Interactions in Ribonucleotide Reductases. Biochemistry 47, 30 (2008).

44. Crona, M. et al. NrdH-Redoxin Protein Mediates High Enzyme Activity in Manganese-reconstituted Ribonucleotide Reductase from Bacillus anthracis. J. Biol. Chem. 286, 33053 (2011).

45. Lin, Q. et al. Glutamate 52-β at theα/β subunit interface of Escherichia coli class Ia ribonucleotide reductase is essential for conformational gating of radical transfer. Journal of Biological Chemistry 292, 9229–9239 (2017).

46. Hasan, M., Banerjee, I., Rozman Grinberg, I., Sjöberg, B. M. & Logan, D. T. Solution Structure of the dATP-Inactivated Class I Ribonucleotide Reductase From Leeuwenhoekiella blandensis by SAXS and Cryo-Electron Microscopy. Front. Mol. Biosci. 8, 1–15 (2021).

47. Minnihan, E. C. et al. Generation of a stable, aminotyrosyl radical-induced α2β2 complex of Escherichia coli class Ia ribonucleotide reductase. Proceedings of the National Academy of Sciences of the United States of America vol. 110 3835–3840 (2013).

48. Lodmell, J. S. & Hennelly, S. P. Conformational Dynamics within the Ribosome. Madame Curie Bioscience Database (Landes Bioscience) (2013).

49. Sharma, S., Luo, M., Patel, H., Mueller, D. M. & Liao, M. Conformational ensemble of Yeast ATP Synthase at Low pH Reveals Unique Intermediates and Plasticity in F1-Fo Coupling. Nat. Struct. Mol. Biol. 31, 657 (2024).

50. Yoshida, M., Muneyuki, E. & Hisabori, T. ATP synthase — a marvellous rotary engine of the cell. Nature Reviews Molecular Cell Biology 2001 2:9 2, 669–677 (2001).

51. Caldwell, C. C. & Spies, M. Dynamic Elements of Replication Protein A at the Crossroads of DNA Replication, Recombination, and Repair. Crit. Rev. Biochem. Mol. Biol. 55, 482 (2020).

52. Doublié, S., Sawaya, M. R. & Ellenberger, T. An open and closed case for all polymerases. Structure 7, R31–R35 (1999).

53. Hammel, M. et al. Ku and DNA-dependent Protein Kinase Dynamic Conformations and Assembly Regulate DNA Binding and the Initial Non-homologous End Joining Complex. J. Biol. Chem. 285, 1414 (2009).

54. Geronimo, I., Vidossich, P. & De Vivo, M. Local Structural Dynamics at the Metal-Centered Catalytic Site of Polymerases is Critical for Fidelity. ACS Catal. 11, 14110–14121 (2021).

55. Strycharska, M. S. et al. Nucleotide and Partner-Protein Control of Bacterial Replicative Helicase Structure and Function. Mol. Cell 52, 844–854 (2013).

56. Blattner, F. R. et al. The Complete Genome Sequence of Escherichia coli K-12. 277(5331), 1453–62 (1997).

57. Lycksell, P.-O. & Sahlin, M. Demonstration of segmental mobility in the functionally essential carboxyl terminal part of ribonucleotide reductase protein R2 from Escherichia coli. FEBS 15778 FEBS Letters 368, 441444 (1995).

58. Sanchez-Garcia, R. et al. DeepEMhancer: a deep learning solution for cryo-EM volume post-processing. Communications Biology 2021 4:1 4, 1–8 (2021).

59. Lopéz-Blanco, J. R. & Chacón, P. iMODFIT: Efficient and robust flexible fitting based on vibrational analysis in internal coordinates. J. Struct. Biol. 184, 261–270 (2013).

60. Kidmose, R. T. et al. Namdinator - automatic molecular dynamics flexible fitting of structural models into cryo-EM and crystallography experimental maps. IUCrJ 6, 526–531 (2019).

61. Chen, V. B. et al. MolProbity: all-atom structure validation for macromolecular crystallography. Acta Crystallogr. D Biol. Crystallogr. 66, 12 (2010).

62. The PyMOL Molecular Graphics System, Version 1.2r3pre, Schrödinger, LLC. - Google Search.

63. Pettersen, E. F. et al. UCSF Chimera--a visualization system for exploratory research and analysis. J. Comput. Chem. 25, 1605–1612 (2004).

