## Supplementary pdf for "CryoEM reveals the dynamic conformational landscape and C-terminal gating mechanism of Mycobacterial Class Ib ribonucleotide reductase"

### 1. Materials and methods:

#### 1.1. Cloning of *nrdE* and *nrdF2* genes from *Mycobacterium thermoresistibile* species:

Mth *nrdE* was cloned in the pGEX4T-3 vector (GST tag) with an ampicillin resistance marker, while the *nrdF2* gene was cloned in the pET28a+ vector (His tag) with kanamycin as a resistance marker. To aid in co-expression and co-purification, different resistance markers, tags and protease site was employed to obtain pure protein complexes.

#### 1.2. Purification of Mth $\alpha_2\beta_2$ complex:

Constructs of the NrdE (α) subunit and NrdF2 (β) subunit were co-transformed in the *BL21 Origami2 (DE3)* strain. 2.5 litre of culture pellet was resuspended in 100 ml lysis buffer containing 50mM Tris pH 7.8, 300mM NaCl, 0.5mM-EDTA, and 3% glycerol. Cells were lysed with the medium probe at 40 amplitude for 2 sec On and 15 sec Off cycle for 1.30 min. The soluble fraction post centrifugation was affinity purified using Glutathione resin for 2 hours at 4ºC. To remove the unbound fractions, washing with 15-column volume of lysis buffer was performed. To obtain native protein, on-column cleavage with thrombin protease was performed to remove the tag. The eluted fraction was concentrated up to 2.0 ml with a 10 kDa Amicon filter. The concentrated protein was injected in the 16/600 Superdex 200 column. The buffer used for SEC chromatography was 25mM Tris pH 8, 150mM NaCl. The homogenous native purified protein obtained was concentrated and used for further experiments.

Reduction of NrdE is essential for the catalytic activity of enzyme. However, in this study our aim was to determine the structure of the complex directly from the native bacterial preparation; therefore, we did not add any reducing agents or nucleotides during purification or prior to grid preparation. Importantly, the resulting structure was similar to that obtained from samples prepared under reducing conditions with nucleotide addition. Nonetheless, we plan to explore alternative solution conditions in future structural studies.

#### 1.3. Glutaraldehyde cross-linking:

To prevent complex dissociation on the grid, the $\alpha_2\beta_2$ complex, purified in a HEPES buffer, was crosslinked with glutaraldehyde for cryo-EM grid preparation. The glutaraldehyde crosslinking was optimised for the time and concentration of glutaraldehyde to be used and evaluated in SDS-PAGE. For grid preparation, $\alpha_2\beta_2$ complex was cross-linked with 0.2% glutaraldehyde for five minutes, and the reaction was terminated using 50 mM Tris buffer at pH 8. Conformer B and conformer D were obtained from crosslinked samples. In contrast, other conformers were obtained from data acquired at CM01 beamline ESRF, Grenoble, France [1], which was not crosslinked and had particles distributed between #10K-#17K.

#### 1.4. Surface plasmon resonance (SPR):

The thermodynamic parameters of α subunit and β subunit interactions were determined using the Biacore T200 system using biosensor analysis at different temperatures from 10-30ºC at an interval of 4 ºC. 800 nM of β subunit protein was immobilised on Sensor chip NTA and 300 nM α subunit as an analyte were flowed over at a rate of 20 µL/min. The buffer used for the SPR experiment was 25 mM Tris buffer (pH 8.0), 300 mM NaCl, 30 µM $MnCl_2$, and 0.02% Tween20, and a similar buffer was used for protein stock solution dilution along with 15 µM TCEP. The thermodynamic parameters were determined using BiacoreT200 evaluation software 3.2. The values of entropy change (ΔS) and enthalpy change (ΔH) were measured by the Van't Hoff equation in linear fit. The Gibbs free energy changes (ΔG) were calculated from the KD values according to Eq $\Delta G=-RT\ln KD$ where T is temperature, R is the universal gas constant, and KD is the equilibrium dissociation constant of the $\alpha_2\beta_2$ complex. The KD value was obtained by fitting data to a 1:1 binding model at different temperatures.

### 2. SUPPLEMENTARY RESULTS:

#### 2.1. Cryo-EM structure determination of $\alpha_2\beta_2$ binary complex

Pairwise sequence alignment analysis of the NrdE, NrdF, and NrdI proteins from *Mycobacterium tuberculosis* (Mtb) and *Mycobacterium thermoresistibile* (Mth) showed sequence identities of 90%, 92%, and 86%, respectively. Due to this high sequence similarity, *M. thermoresistibile* was used as a substitute, as we encountered difficulties in purifying the Mtb proteins

When the β subunit is co-expressed with the GST affinity tagged α subunit, it can be clearly seen that the two proteins co-elute on size exclusion chromatography **(Supplementary Figure 2B).** The intact binary complex obtained from cryo-EM data was subjected to 5 classes of heterogeneous refinement since conformational and compositional heterogeneity was observed **(Supplementary Figure 3).** Since, particles from Class 1 and Class 2 were very similar, they were merged and subjected to homogenous refinement, followed by global CTF and non-uniform refinement **(Supplementary Figure 3C)** [2]. This merged particle is called as conformer D. Class 3 particles (named conformer B), were utilized for low-resolution model building through homogeneous refinement. Local refinement techniques were subsequently employed to enhance the reconstruction resolution **(Supplementary Figure 3D).** Due to insufficient numbers of particles and poor map, Class 4 and Class 5 particles were discarded **(Supplementary Figure 3).** The maps of conformers A, C, G, E, F and G were obtained from different non-crosslinked ternary complex data. Since the particles are divided into various classes, the overall resolution of the map here also is compromised **(Supplementary Figure 1).**

#### 2.2. Global conformational changes across conformers:

Broad variations in assembly observed in the α and β subunits of α2β2 complex across all seven conformers were assessed by calculating the center of mass (COM) for each of the four chains **(Supplementary Figure 4A-4G).** Similarly, the same was compared with the COM measurements in the *S. typhimurium* and *E. coli* structures **(Supplementary Figures 4H and 4I).** In all the seven different conformers, minor variations in COM distance are observed between two monomers of the α subunit and between two monomers of the β subunit, suggesting minimal mobility within each dimer. However, variations between α and β, α and β', α' and β, and α' and β', show larger fluctuations ranging upto 8Å (α-β), 9Å (α-β'), 18Å (α'-β), and 25Å (α'-β') **(Supplementary Figures 4A-4G).** Similarly, pronounced variability is observed in the angles formed between the center of mass of α'αβ (66-85°), α'βα (51-69°), βα'α (42-45°), and ββ'α' (19-62°), indicating significant changes in the association of the subunits.

More detailed analysis of inter-subunit variations was undertaken by first superposing the α chain to bring them into one single frame of reference for all the seven conformers (and that of *S. typhimurium*) and then calculating transformations required to superpose other α' chain with respect to this frame of reference. Comparison of the *S. typhimurium* structure with all seven conformers shows rotations in the range of 22- 33° for α' chain (Supplementary Table 1). On the other hand, comparisons within the seven conformers of Mth RNR show the same rotations in the range of 2- 20° (Supplementary Table 1). Similar analysis with β chain as a frame of reference shows 16- 30° rotation of the β' chain when seven conformers are compared with *S. typhimurium* structure, but only 3.5- 19° rotational variation is observed within the seven conformers **(Supplementary Table 2).**

**2.3. Insights from in-solution studies reveal the dynamic nature and flexibility in the $\alpha_2\beta_2$ complex:**

Thermodynamic analysis using SPR reveal conformational flexibility present in-solution in the RNR complex. Surface plasmon resonance analysis confirms that the interaction between α and β subunits is enthalpically driven **(Supplementary Figure 9).** The binding kinetics exhibited notable temperature dependence, indicating that the interaction involves the stabilization of flexible regions upon complex formation. As the temperature increased, the affinity between the α and the β subunits decreased, suggesting that the interaction becomes less favourable at higher temperatures **(Supplementary Figure 9A).** This finding is also supported by the cryo-EM structure, which revealed the presence of flexible C-terminal and N-terminal tails at the interaction interface, which plays a crucial role in mediating the interaction between the α and the β subunit and thereby facilitating radical transfer. These regions become more ordered upon binding, contributing to the enthalpic stabilization of the complex, albeit with unfavourable entropic changes. The observed unfavourable entropic changes during the interaction of the α and the β subunits suggest a decrease in the system disorder. This implies that the molecules are organizing or arranging themselves in a more structured manner as the reaction progresses.

**References:**

1. Kandiah, E. *et al.* CM01: a facility for cryo-electron microscopy at the European

Synchrotron. *Acta Crystallogr. Sect. D, Struct. Biol.* **75**, 528 (2019).

2. Punjani, A., Zhang, H. & Fleet, D. J. Non-uniform refinement: adaptive regularization improves single-particle cryo-EM reconstruction. *Nat. Methods 2020 1712* **17**, 1214–1221 (2020).

### Supplementary Figures:

### Supplementary Figure 1

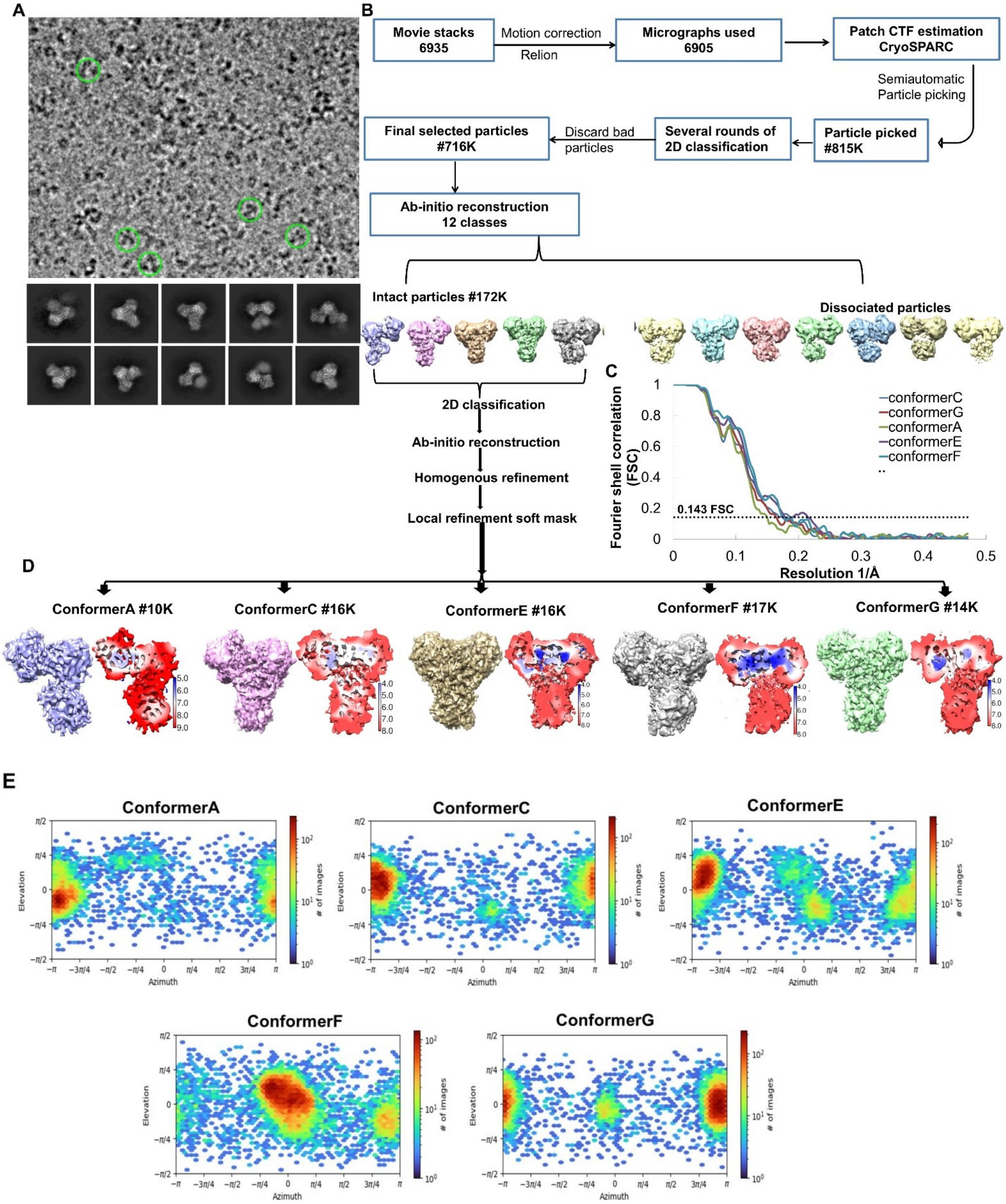

**Supplementary Figure 1: Single-particle cryo-EM processing workflow and reconstructions of the Mth NrdEFI complex**

**A.** Micrograph highlighted with particles and 2D class averages of the ternary complex. **B**. Workflow of cryo-EM data processing for Mth NrdEFI complex. **C.** Gold standard Fourier shell correlation curve for different conformations. The dashed line represents the overall nominal resolution of each reconstruction at 0.143 of FSC calculated using CryoSPARC. The resolution range and resolution of the map are mentioned in Table 1. **D.** Cryo-EM map of different conformational states along with local resolution estimation with the color scale bar shown on its left indicates resolution in Å. **E.** Angular distribution plot for different conformers generated in cryoSPARC showing particle projections. The heat map shows the number of particles corresponding to each viewing angle.

**Supplementary Figure 2**

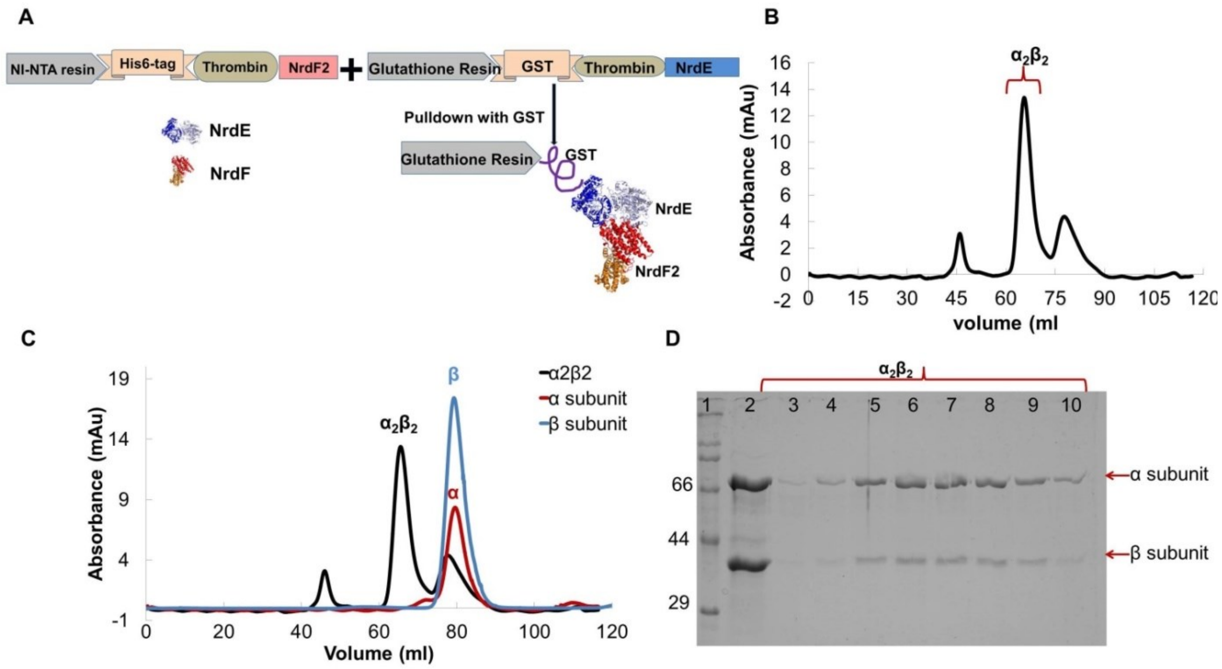


**Supplementary Figure 2: Purification of Mth $\alpha_2\beta_2$ complex**

**A.** The cartoon represents the details of the constructs ($His_6$ tagged NrdF2/β subunit and GST tagged NrdE/ α subunit) used for co-expression, pull-down, and co-purification of the complex. **B.** Size exclusion chromatography profile of affinity purified fraction shows homogenous heterotetrameric complex. **C.** Size exclusion chromatography profiles for β subunit (blue), α subunit (red), and the $\alpha_2\beta_2$ (black) complex overlaid to highlight their respective elution volumes. **D.** 10% SDS gel analysis of Mth $\alpha_2\beta_2$ complex after SEC purification. Lane 1- Molecular weight marker, 2- Sample injected in SEC, 3-10 SEC purified fractions loaded for the peak highlighted with curly bracket in B.

### Supplementary Figure 3

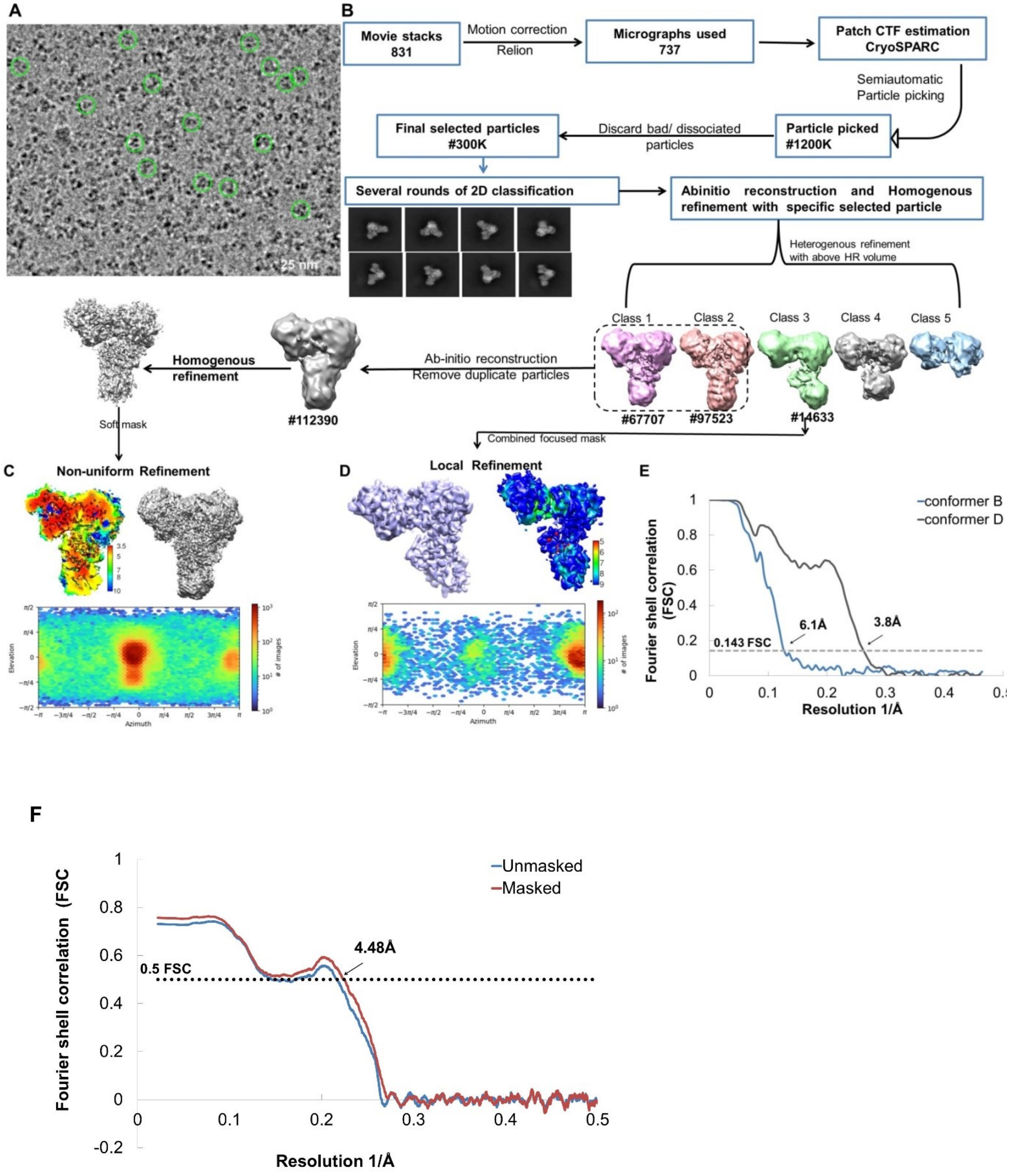

**Supplementary Figure 3: Single-particle cryo-EM processing workflow and reconstructions of the Mth $\alpha_2\beta_2$ complex**

**A.** Micrograph highlighted with particles of $\alpha_2\beta_2$ complex. **B.** Workflow of cryo-EM data processing for $\alpha_2\beta_2$ complex**.** **C.** Conformer D: Local resolution estimation with the color scale bar shown on its right indicates resolution in Å, Cryo-EM map of conformer D of $\alpha_2\beta_2$ complex, Angular distribution heat map of particle projections. **D.** Conformer B: Cryo-EM map of conformer B, Local resolution estimation, and Angular distribution heat map of particle projections. **E.** Gold standard Fourier shell correlation curve for conformer D and conformer B. Dashed line represents the overall nominal resolution of each reconstruction at 0.143 FSC calculated by CryoSPARC. **F.** Map model FSC curve of conformer D. Dashed line represents the resolution of each reconstruction at 0.5 FSC obtained from PHENIX

**Supplementary Figure 4**

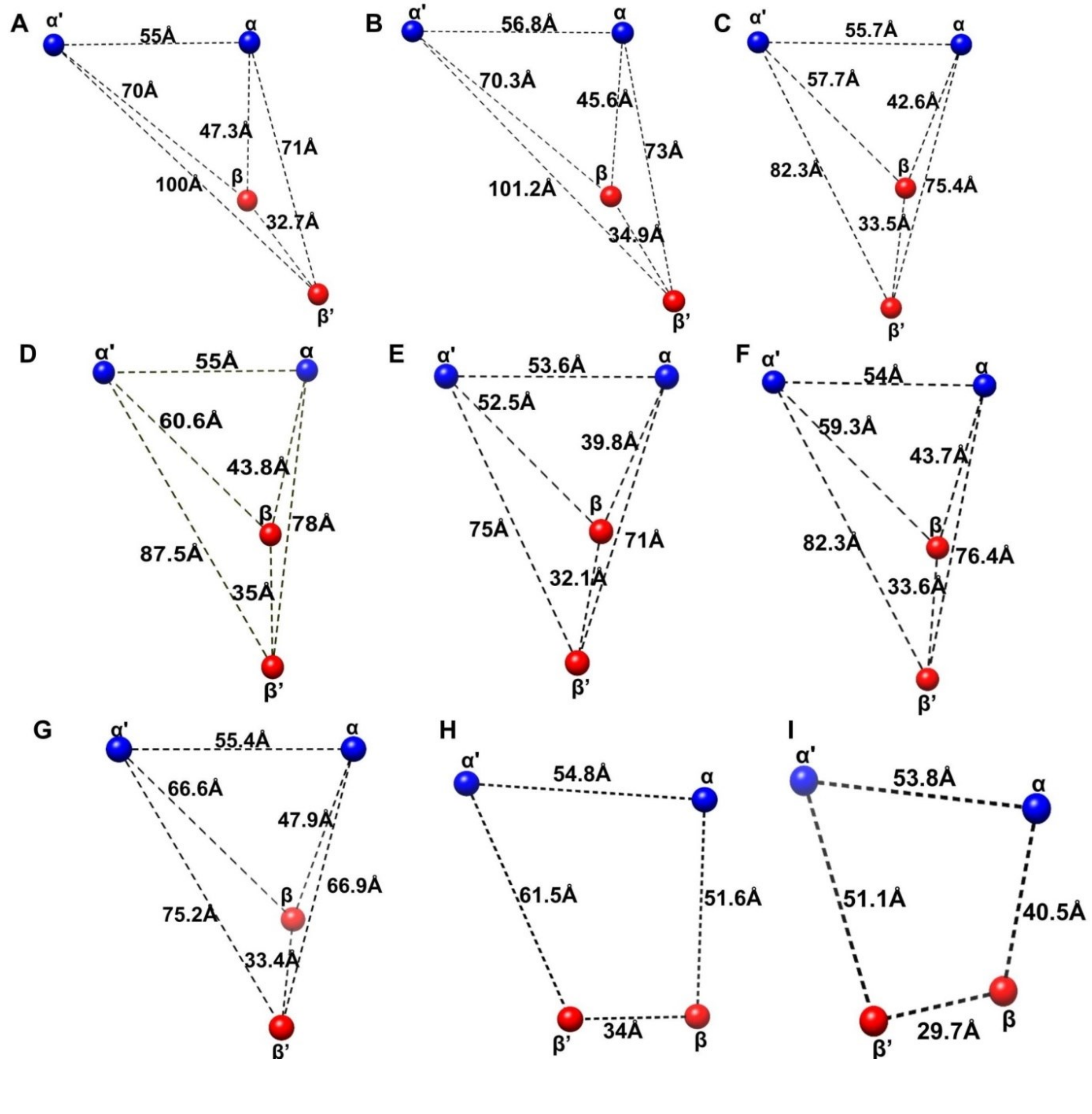


**Supplementary Figure 4: Global conformational change in $\alpha_2\beta_2$ complex**

The global conformational change observed in α and β subunits in the RNR complex. **A-G.** Centre of Mass analysis for Conformer A- Conformer G, **H.** COM for *S. typhimurium* structure, **I.** COM for *E. coli* structure.

**Supplementary Figure 5**

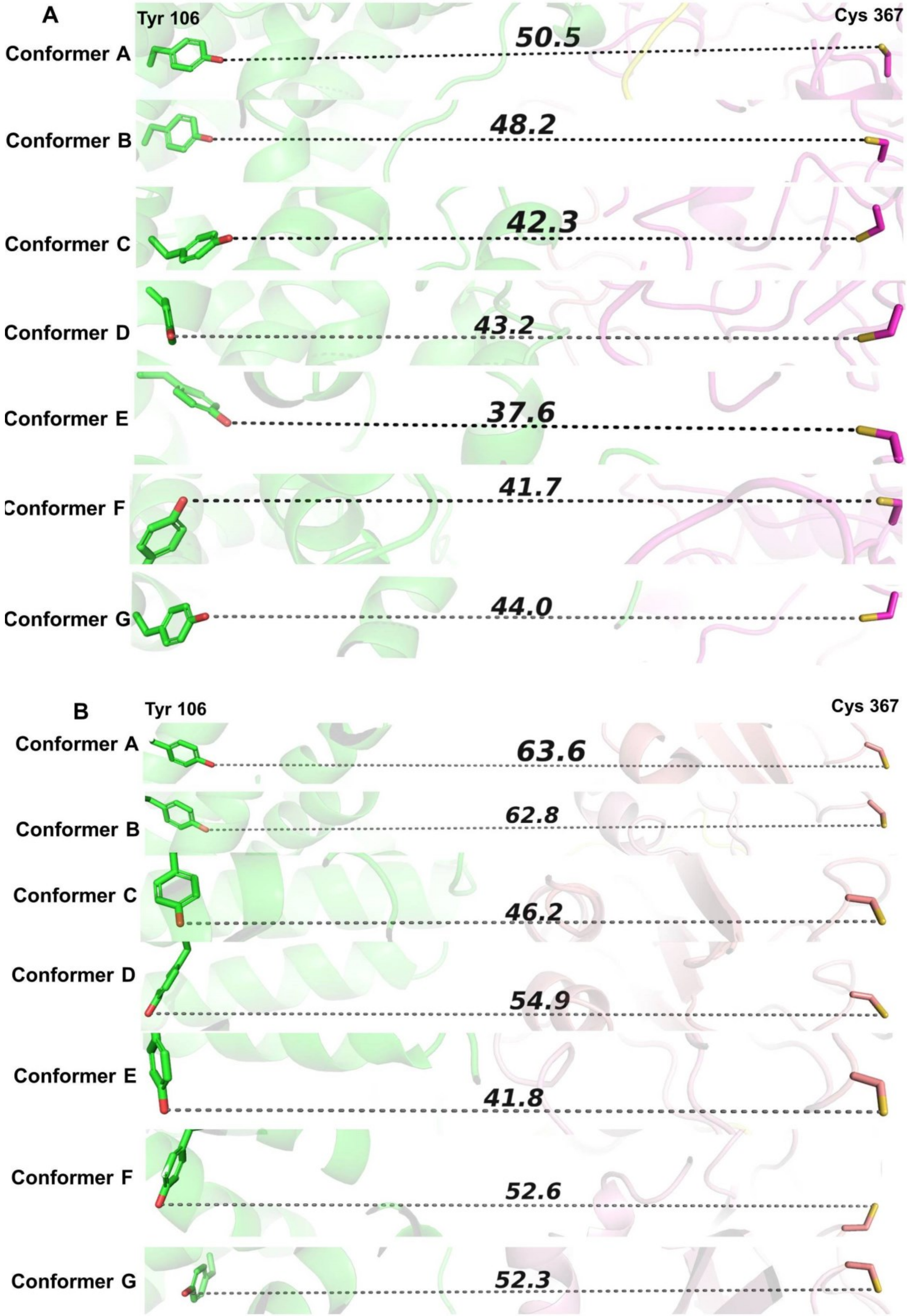

**Supplementary Figure 5: Radical transfer distance from α subunit to β subunit**

The different conformation of the $\alpha_2\beta_2$ complex demonstrates that the flexibility of the β subunit in the complex leads to variable radical transfer distance. **A.** Distance from active site cysteine (367) of α chain to metal cofactor site tyrosine (106) of β chain. **B.** Distance from active site cysteine of α' chain to metal cofactor site tyrosine of β chain. The distance is shown in Å for the different conformers as labelled.

**Supplementary Figure 6**

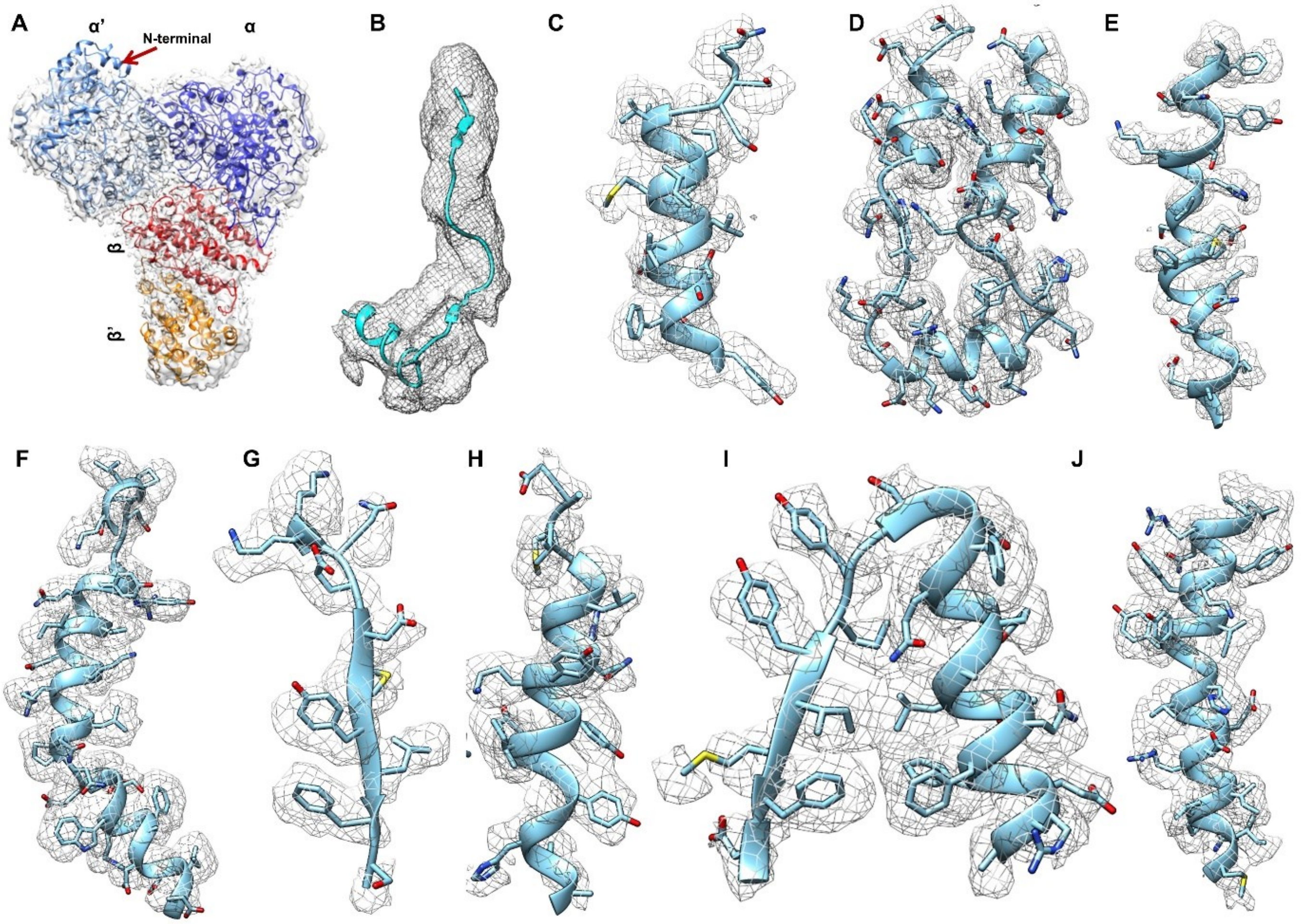


**Supplementary Figure 6: Map model superposition of $\alpha_2\beta_2$ complex**

**A.** Map (Unsharpened map) - model superposition of $\alpha_2\beta_2$ complex. The arrow indicates the region where the density for the N-terminal region is missing. Cryo-EM map demonstrating details in different regions of the map. **B.** Unsharpened map highlighting C terminal tail region of β chain. CryoSPARC sharpened map demonstrating density from different regions and chains of complex, **C.** chain E (aa 206-223), **D**. chain F (aa 509-553), **E.** chain A (aa 88-110), **F**. chain B (aa 118-150), **G**. chain F (aa 280-289). **H.** chain A (aa 194-212), **I.** chain E (aa 326-348), **J.** chain B (aa 185-208). The map was contoured at 0.1-0.15σ to provide completeness in mapping density around the atomic coordinates. Colour zone tool of chimera with radius cutoff of 2Å was used to map the density in different chain of conformer D.

**Supplementary Figure 7**

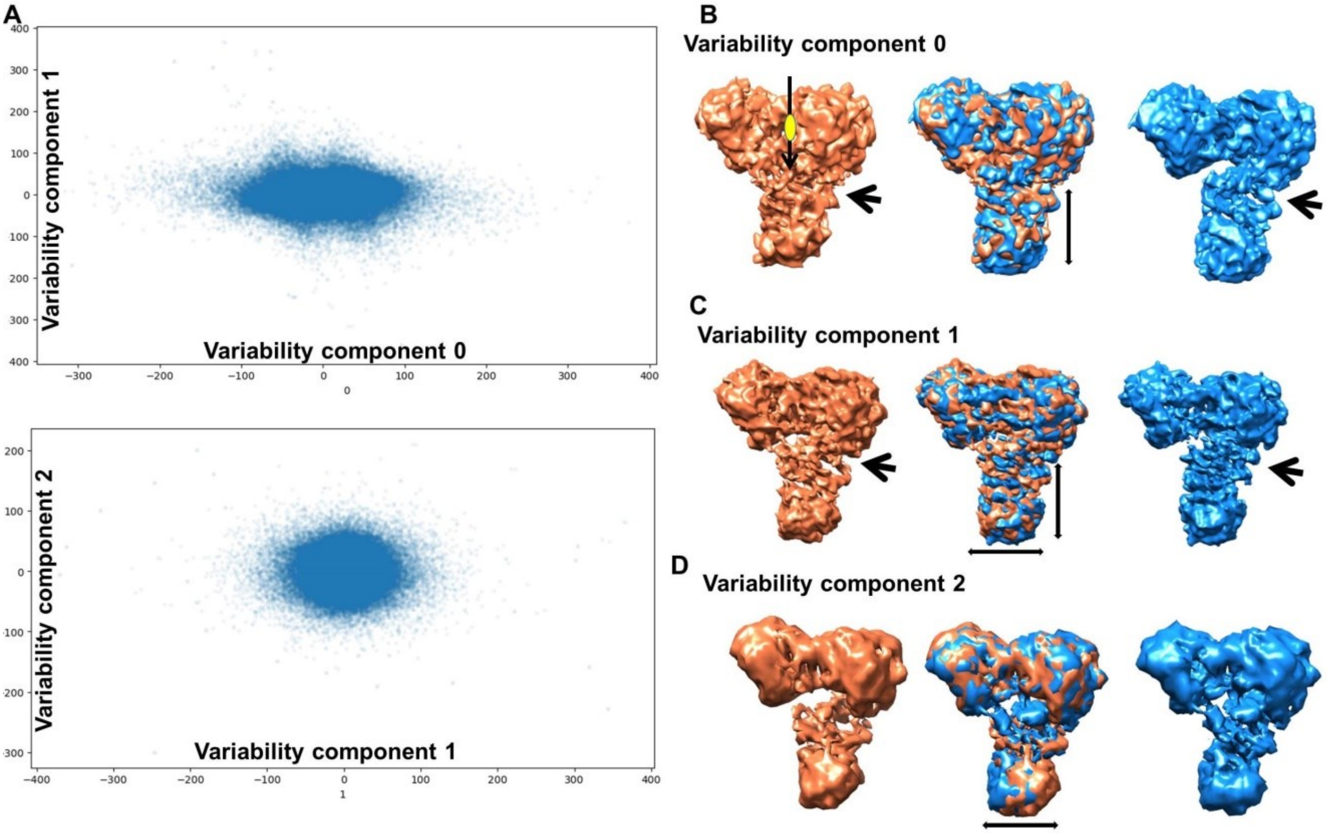


**Supplementary Figure 7: Variability analysis of $\alpha_2\beta_2$ complex**

3D variability analysis of conformer D. Scatter plot obtained for variability along **A.** Component 0 and 1 and components 1 and 2 present in the sample. **B.** Variability Component 0. A yellow ellipsoid indicates the identified 2-fold axis. **C.** Variability Component 1 Arrow indicates variability observed near the C-terminal region of the β chain. **D.** Variability Component 3. Movement along the mean position is indicated by a double-headed arrow.

**Supplementary Figure 8**

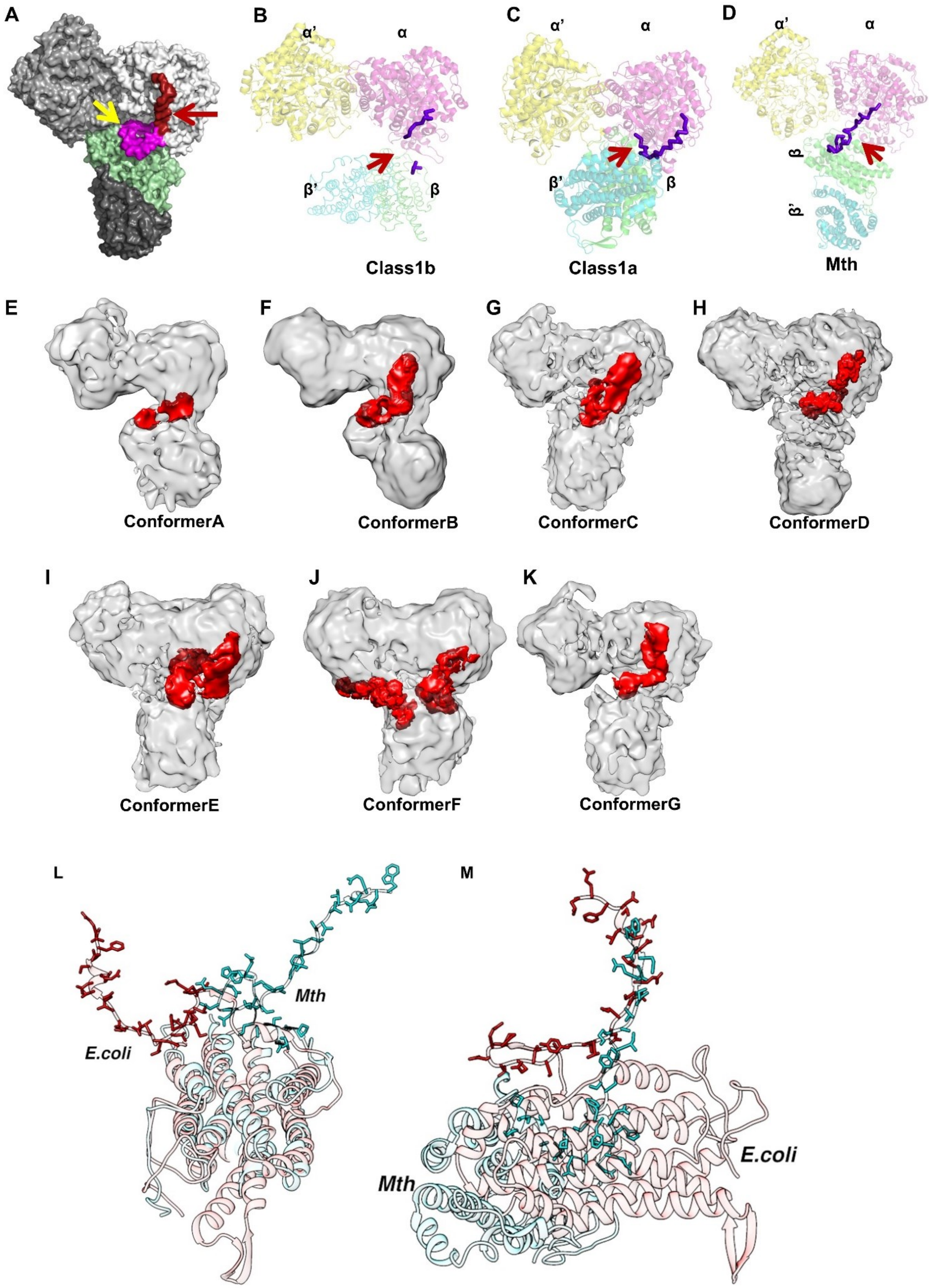

**Supplementary Figure 8: C-terminal tail interaction of β chain of RNR** **A.** Surface representation of Mth $\alpha_2\beta_2$ complex of conformer D revealing the first half magenta (indicated by yellow arrow) and second half red (indicated by red arrow) C-terminal tail of β chain. **B.** *S.typhimurium* class Ib structure with a missing portion of C-terminal indicated by red arrow. **C.** *E. coli* class Ia doubly substituted trapped structure with continuous C-terminal in purple indicated by red arrow. **D.** Cartoon representation of the Mth $\alpha_2\beta_2$ structure, with the C-terminal tail highlighted in purple. **E-G.** Unsharpened map of different conformers (conformer A to conformer G) with the region of the C-terminal tails highlighted in red. **L-M.** Superposition *E.coli* (6W4X shown in red) and Mth (conformer D shown in cyan) structure to demonstrate the orientation of C-terminal tail in class Ia and class Ib RNR. **L**. Superposition of β-subunit of both structures and **M.** Superposition of α-subunit demonstrating variation observed in the C-terminus of β-subunit.

**Supplementary Figure 9**

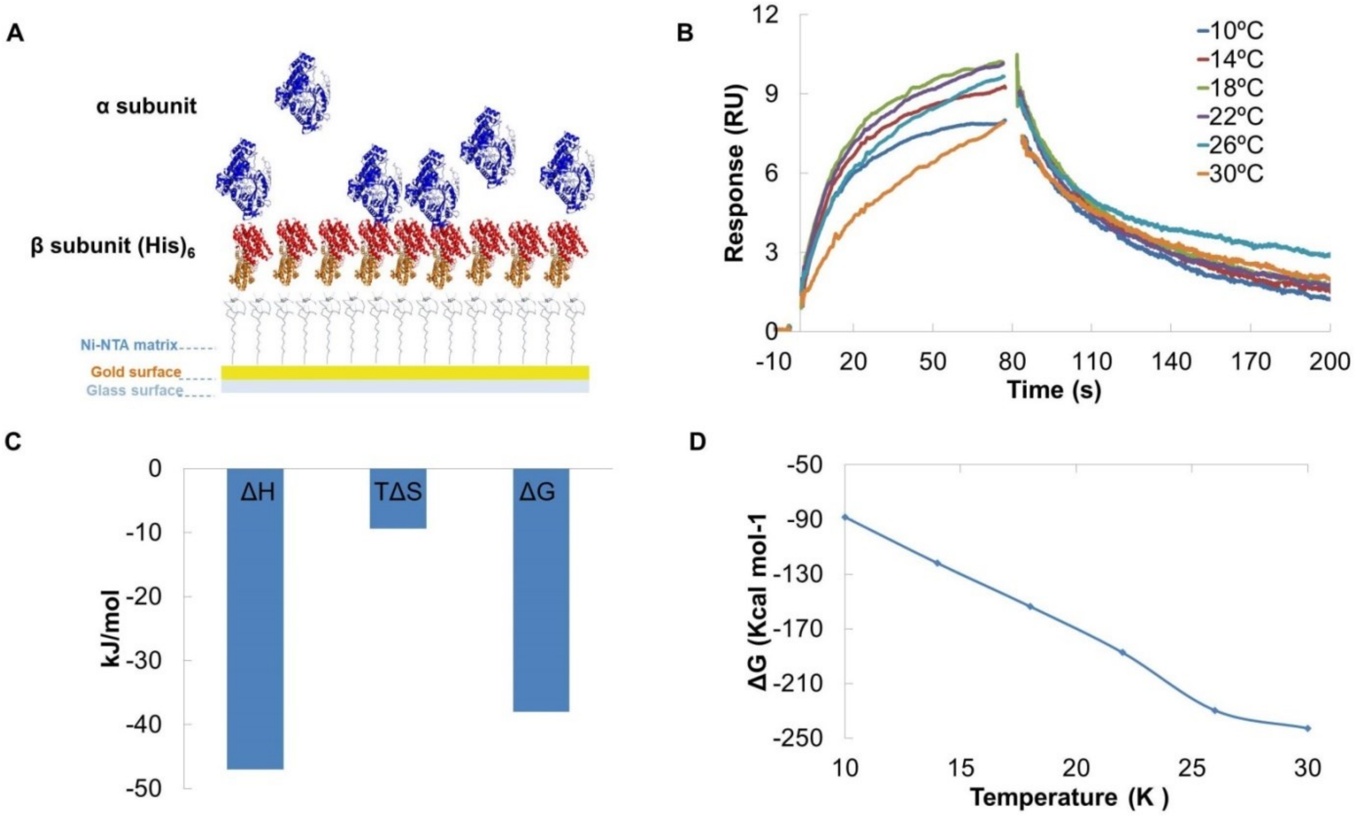


**Supplementary Figure 9: The effect of temperature on kinetics of α subunit interaction on immobilized β subunit**

**A.** Cartoon representation of SPR reaction set up for α subunit interaction on Immobilized β subunit using Ni-NTA matrix. **B.** Plot of $\alpha_2\beta_2$ interaction sensorgram at different temperatures ranging from 10-30°C at an interval of 4°C. The colors of the sensorgram are indicated as 10°C-blue, 14°C-red, 18°C- green, 22°C- violet, 26°C- light blue, 30°C-orange. **C.** Bar graph showing enthalpy and entropic contribution to the free energy of binding of $\alpha_2\beta_2$ interaction. **D.** Change in Gibbs free energy of binding at different temperatures from 10-30°C at an interval of 4°C.

**Supplementary Figure 10**

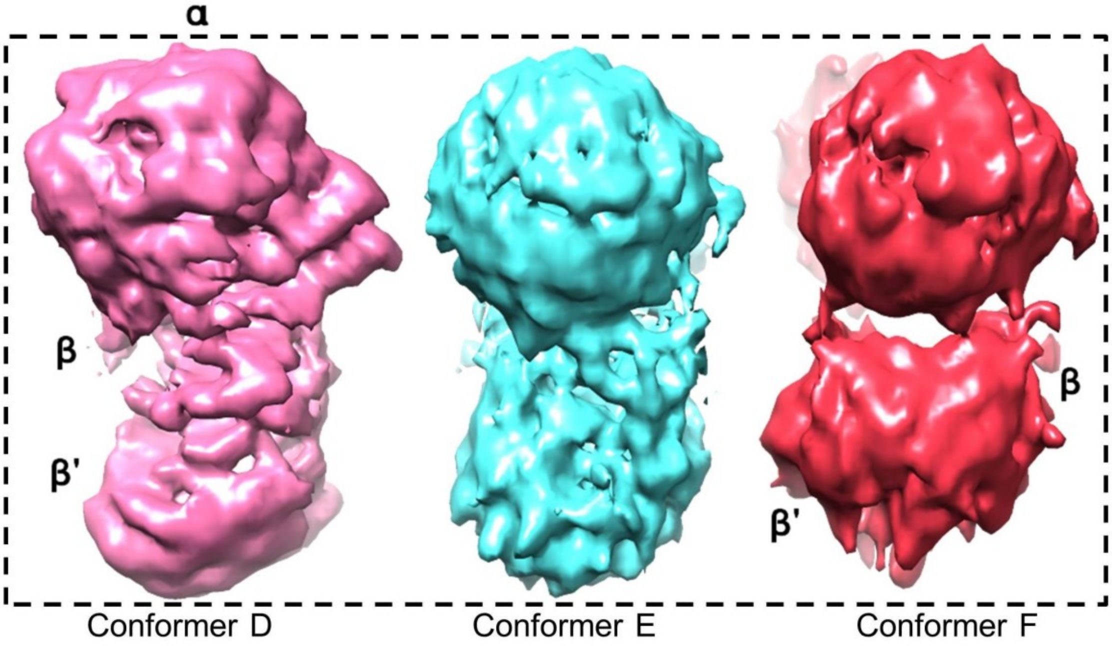


**Supplementary Figure 10: Compaction and bending in β and β' chain**

Unsharpened cryoEM map of conformers D, E, and F demonstrating bending in α, β, and β', compaction in conformer E leading to a decrease in radical transfer distance, and bending and compaction in β and β' chain in conformer F with a reduction in radical transfer distance promotes closer interaction between α' and β'.

**Supplementary Tables:**

**Supplementary Table 1**

| | Conformer G | Conformer E | Conformer A | Conformer F | Conformer C | Conformer B | Conformer D |
|---|---|---|---|---|---|---|---|
| 2BQ1 | 22 | 26 | 33 | 24.9 | 27.2 | 26 | 29.9 |
| Conformer G | | 6.2 | 13.6 | 2.7 | 9 | 8.8 | 3.9 |
| Conformer E | 6.2 | | 17 | 5.7 | 13.2 | 10.9 | 7.6 |
| Conformer A | 13.6 | 20 | | 15.5 | 13.7 | 9.11 | 16 |
| Conformer F | 2.7 | 5.7 | 15.5 | | 8.7 | 8.4 | 1.7 |
| Conformer C | 9.6 | 13.2 | 13.7 | 8.7 | | 6 | 8.6 |
| Conformer B | 8.8 | 11 | 9 | 8.4 | 6 | | 9.8 |
| Conformer D | 3.9 | 7.6 | 16 | 1.7 | 8.6 | 9.8 | |

**Supplementary Table 1: Least square fit analysis of α subunit:**
The rotation angle was calculated for α' chain among the seven conformers and *S. typhimurium* keeping (α chain) as fixed frame of reference.

**Supplementary Table 2**

| | Conformer A | Conformer B | Conformer C | Conformer D | Conformer E | Conformer F | Conformer G |
|---|---|---|---|---|---|---|---|
| 2BQ1 | 21.4 | 30 | 24 | 26.7 | 27 | 16 | 23 |
| Conformer A | | 7.2 | 13.5 | 14 | 4 | 8.2 | 15.4 |
| Conformer B | 7.2 | | 11.6 | 9.8 | 5.8 | 18.9 | 11.6 |
| Conformer C | 13.5 | 11.6 | - | 4.4 | 9.6 | 15.4 | 4 |
| Conformer D | 14 | 9.8 | 4.4 | - | 8.7 | 18.7 | 3.3 |
| Conformer E | 4 | 5.8 | 9.6 | 8.7 | - | 12 | 10.9 |
| Conformer F | 8.2 | 18.9 | 15.4 | 18.7 | 12 | - | 16.5 |
| Conformer G | 15.4 | 11.6 | 4 | 3.3 | 10.9 | 16.5 | - |

**Supplementary Table 2: Least square fit analysis of β subunit:**
The rotation angle was calculated for β' chain among the seven conformers and *S. typhimurium* keeping (β chain) as fixed frame of reference.

**Supplementary Table 3**

| | Conformer A | | Conformer B | | Conformer C | | Conformer E | | Conformer F | | Conformer G | |
|---|---|---|---|---|---|---|---|---|---|---|---|---|
| | β | β' | β | β' | β | β' | β | β' | β | β' | β | β' |
| Conformer D | 46 | 45 | 37.8 | 44.3 | 1.4 | 5 | 5.8 | 9 | 44.6 | 62.6 | 1.69 | 2.14 |

**Supplementary Table 3: Least square fit analysis of β relative to α:**
The table shows the rotation angle calculated for β and β' chain relative to α chain among all seven conformers.

### Supplementary Table 4

| | | | | Interface area (Å²) | | | | | | |
|---|---|---|---|---|---|---|---|---|---|---|
| | | | | Conformer | | | | | | |
| Chain ID | 6W4X | 2BQ1 | 9BZ9 | A | B | C | D | E | F | G |
| α:α’ | 1351.8 | 1166.7 | 1696.6 | 1329.2 | 1374 | 1507.3 | 1549.4 | 1212 | 1697.5 | 1550.6 |
| β:β’ | 3344.4 | 1668.3 | 1048.5 | 1494 | 1678.6 | 1536.2 | 1363.1 | 2208 | 1415.7 | 1317.7 |
| α:β | 3352.3 | 1238.5 | 1025 | 962.7 | 1175.7 | 1649.1 | 1347.3 | 2094 | 824.9 | 2048.2 |
| α’:β’ | 500 | - | 814.3 | - | - | - | 223.4 | - | 461.1 | - |
| α’β | - | - | 90 | - | - | 421.8 | - | 580.0 | 49.7 | 688.2 |

**Supplementary Table 4: Interface area of $\alpha_2\beta_2$ complex:**
The interface area in Å² of $\alpha_2\beta_2$ complex structure of *E. coli*, *S. typhimurium, B. subtilis* and Mth seven conformers were assessed using PDBePISA. The interface area between the dimer of the α subunit, dimer of β subunit, α and β chain, α’ and β chain, and α’ and β’ chain is shown in the table.

### Supplementary Table 5

| Data collected | α2β2 sample | EFI sample |
|---|---|---|
| #of images collected | 831 | 6935 |
| Magnification (x) | 130000 | 130000 |
| No. of frames | 32 | 40 |
| Dose rate (e-/p/s) | 6.8 | 9.3 |
| Exposure time (s) | 8 | 5 |
| Pixel size (Å) | 1.065 | 1.052 |
| Dose/frame (e-/Å2) | 1.48 | 1.05 |
| Total Dose (e-/Å2) | 47.96 | 42.02 |
| Objective aperture (ìm) | 100 | 100 |
| Defocus range (ìm) | 1.5-3.5 | 0.7 - 3.5 |

**Supplementary Table 5:** Data collection parameters for $\alpha_2\beta_2$ binary complex and NrdEFI complex

**Supplementary Videos:**

**Supplementary Video 1:** Overview demonstrating different major conformational transitions from conformer A to conformer F from structure determined using cryo-EM map

**Supplementary Video 2:** Close view of the conformational shift of β hairpin loop away from the active site with the docking of the C-terminal tail of the β subunit towards the active site observed on the close interaction of α and β subunit in conformer D.

**Supplementary Video 3:** Continuous heterogeneity of conformer D observed along components 0, 1, and 2. The different component resolves the synchronous motion of α and β subunits, promoting closer interaction with each other

**Supplementary Video 4:** Conformational transition of conformer A and conformer B visualized through morphing using an unsharpened map.

**Supplementary Video 5:** Conformational transition of conformer B and conformer D visualized through morphing using an unsharpened map.

**Supplementary Video 6:** Conformational transition of conformer D and conformer E visualized through morphing using an unsharpened map.

**Supplementary Video 7:** Conformational transition of conformer E and conformer F visualized through morphing using an unsharpened map.

**Supplementary Video 8:** Transition of conformer A to conformer F visualized through morphing, using an unsharpened map to demonstrate the migration of α' and β' subunits towards each other. Two different levels of the map are shown for clarity.

**Supplementary Video 9:** Mechanism of interaction between α and β chains and the hypothesized mode of association between α' and β' chains.
